# CodonMamba: a foundation model for programmable mRNA coding sequence design

**DOI:** 10.64898/2026.08.24.746601

**Authors:** Mei Lang, Xingyu Fang, Zhen Wang, Mingxuan Chen, Zhaowen Cheng, Xiagu Zhu, Kin Yip Tam, Junwei Zhang, Xiaolin Li

## Abstract

Although mRNA codon language models provide a generalizable framework for biological sequence design, effective CDS design requires both a learned sequence design space that captures biological constraints and context-configurable design preferences. Here we present CodonMamba, a codon language model framework for mRNA prediction and programmable CDS design. Pretrained on large-scale coding-sequence corpora, CodonMamba achieved state-of-the-art performance across a comprehensive benchmark of 12 mRNA prediction tasks, ranking first on 10 of 12 tasks. Furthermore, CodonMamba establishes a programmable CDS design framework, transforming codon optimization from a process dependent on preferences embedded during model training into an inference-time steerable generation framework. By introducing user-specified codon usage priors during generation, CodonMamba enables inference-time steering toward host- or application-specific codon preferences without retraining. In design experiments, CodonMamba enabled coordinated optimization over multiple design-relevant sequence properties and demonstrated programmable cross-host CDS retargeting through inference-time prior switching, while preserving most model-derived codon choices.Together, these results establish CodonMamba as a programmable foundation model framework for CDS design, enabling context-specific mRNA sequence engineering and highlighting a promising direction for precise mRNA design using foundation models.

## 1 Introduction

Messenger RNA (mRNA) therapeutics have emerged as a versatile platform for programmable protein delivery, enabling rapid cycles of design, construction and testing without genomic integration [1]. Their clinical potential was established by the success of mRNA vaccines during the COVID-19 pandemic [2, 3] and has since expanded into oncology, where personalized neoantigen vaccines are showing encouraging clinical activity [4–6]. These advances have increased the need for computational approaches that can systematically engineer mRNA sequences for diverse biological settings.

A central challenge in mRNA coding sequence (CDS) engineering is that effective sequence design depends on both biological context and the ability to impose context-specific sequence preferences. Because of the degeneracy of the genetic code, a single protein can be encoded by a vast number of synonymous CDSs that preserve the same amino acid sequence yet differ substantially in translation, stability and cellular compatibility [7]. The preferred sequence properties are not necessarily universal, but can depend on the biological setting in which the mRNA is intended to function, including the expression host and, more broadly, tissue- and cell-specific translational environments [8–11]. Therapeutic CDS design therefore requires more than identifying a generally plausible synonymous sequence: it requires a model that captures broad biological sequence regularities while allowing sequence preferences to be adapted to the intended context.

Traditional CDS optimization methods provide explicit control over predefined sequence objectives. Codon-usage approaches typically represent the target host through species-level codon frequencies or derived measures such as the codon adaptation index (CAI) [12–15], whereas LinearDesign additionally incorporates RNA secondary-structure free energy through dynamic programming [16]. Although efficient and interpretable, such approaches rely on explicitly specified objectives and therefore capture only a limited subset of the higher-order dependencies among synonymous codon choice, nucleotide composition, RNA structure and local sequence context.

Language models provide a complementary strategy by learning rich sequence distributions directly from large-scale and heterogeneous natural data, thereby capturing higher-order dependencies that are difficult to encode explicitly [17– 22]. Recent codon and mRNA language models have demonstrated strong performance in prediction and sequence optimization [23–27]. However, learning a general sequence distribution does not by itself enable sequence preferences to be flexibly reconfigured for different application contexts. In some current frameworks, context-specific sequence preferences are coupled to the training distribution, conditioning scheme or optimization objective, limiting the ability to reconfigure context-specific preferences at inference without additional training [24, 28].

Recent studies in protein design have shown that pretrained generative models can be redirected towards user-specified properties at generation time through activation-based steering or predictor-based guidance [29, 30]. Protein-preserving CDS design provides a particularly interpretable setting for direct inference-time control. Once the target protein is fixed, genetic-code degeneracy defines an explicit set of admissible synonymous codons at each position [7, 14]. Among the sequence properties considered in CDS optimization, host codon-usage preference is a central and widely used design criterion, reflecting systematic differences in synonymous codon usage across expression hosts [12–14]. Relative codon adaptiveness, originally introduced in the codon adaptation index framework [12], provides a quantitative representation of this preference by measuring the relative usage of each codon within its synonymous family. Host-specific codon-usage preferences can therefore be translated into codon-level weights that act directly on synonymous codon selection during constrained decoding. This motivates a framework in which broad coding-sequence regularities learned during pretraining are retained, while protein-preserving constraints define the admissible design space and replaceable application-specific preferences steer synonymous codon selection within that space without modifying the pretrained model parameters.

Here we present CodonMamba (Fig. 1), a codon language model framework for mRNA prediction and programmable CDS design. CodonMamba uses bidirectional state-space blocks to integrate coding-sequence context from both directions [31] and was pretrained on approximately 9 million CDSs from 1,544 organisms using masked codon prediction. As a first test of whether the pretrained backbone captured broadly informative coding-sequence regularities, we evaluated its representations across 12 downstream mRNA prediction benchmarks against 11 pretrained baseline models spanning DNA, RNA and protein modalities. CodonMamba ranked first on 10 tasks and second on the remaining two, demonstrating broad and consistent transferability across heterogeneous mRNA functions.

**Figure 1.**
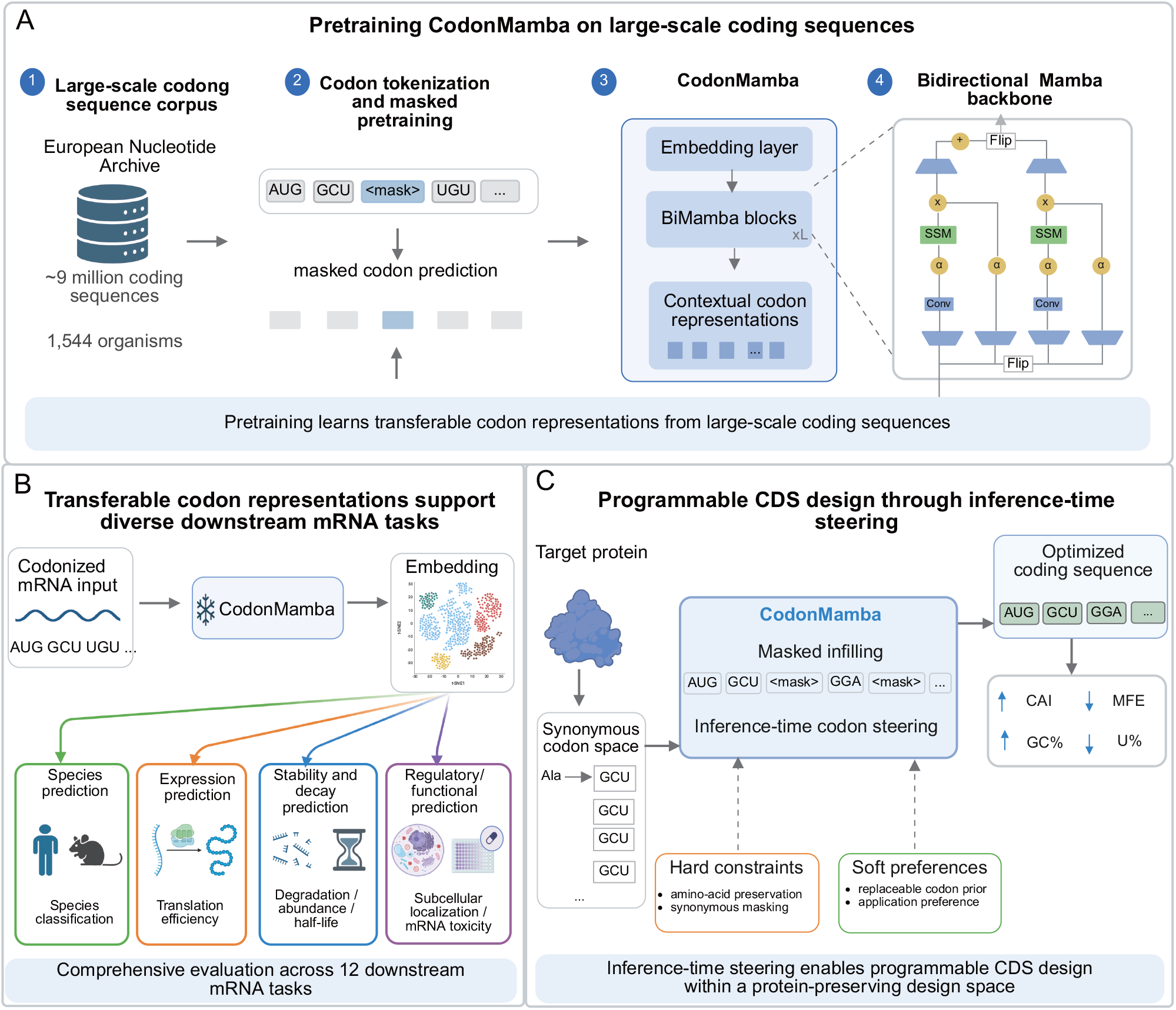
Overview of CodonMamba for coding-sequence modelling and programmable CDS design. **A**, Pretraining corpus and model architecture. CodonMamba was pretrained on approximately 9 million coding sequences from 1,544 organisms using masked codon prediction and a bidirectional Mamba backbone. **B**, Downstream evaluation across 12 mRNA prediction tasks spanning protein expression, mRNA stability and degradation, and regulatory phenotypes. **C**, Constrained CDS generation by iterative masked codon infilling. Synonymous-codon masking restricts decoding to protein-preserving alternatives, whereas a host codon usage prior provides a soft, replaceable inference-time preference over admissible codons.

Building on this broadly transferable coding-sequence representation, CodonMamba implements programmable CDS design through constrained masked infilling. Synonymous-codon masking defines a protein-preserving design space, whereas a user-specified host codon usage prior provides a replaceable inference-time preference without retraining. Switching this prior systematically retargeted generated CDSs towards the codon-usage preferences of four phylogenetically diverse hosts while preserving most model-preferred codon choices. CodonMamba further produced coordinated shifts across multiple design-relevant sequence properties. Together, these results establish a framework that separates learned coding-sequence distributions from context-configurable design, enabling the same pretrained model to accommodate different application-specific sequence preferences through lightweight inference-time steering.

## 2 Results

### 2.1 CodonMamba captures multi-scale coding-sequence regularities through self-supervised pretraining

Because both downstream transfer and programmable decoding depend on the pretrained backbone to provide informative coding-sequence representations, we first asked whether self-supervised pretraining captured established biological regularities across multiple scales. Following representation analyses established in previous codon language models such as CaLM and CodonBERT [24, 32], we examined CodonMamba representations at two complementary levels: static embeddings of the 64 standard codons (Fig. 2A–C) and contextual sequence-level embeddings from an independent held-out species dataset (Fig. 2D–E).

**Figure 2.**
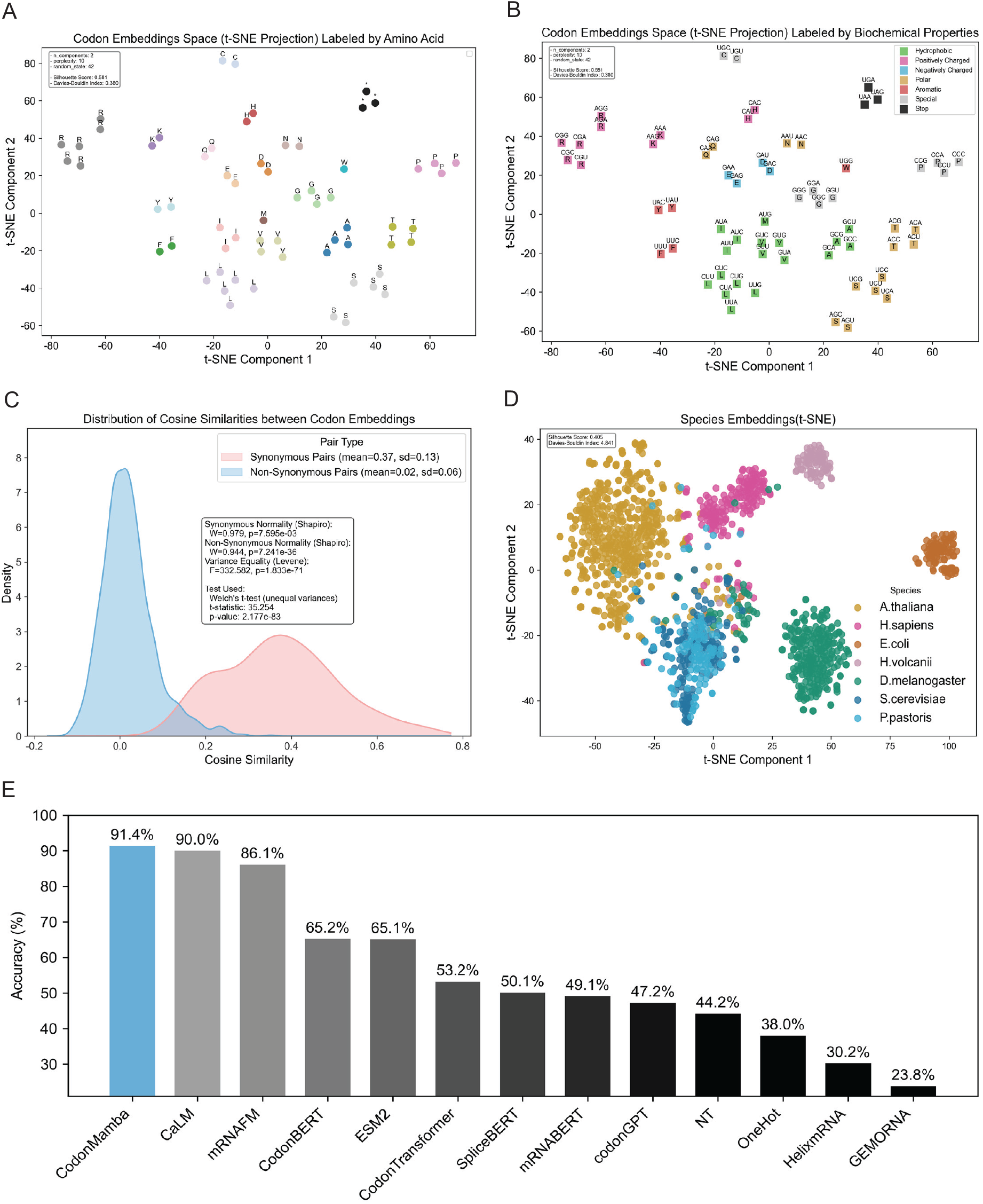
CodonMamba captures multi-scale coding-sequence organization through self-supervised pretraining. **A**, t-SNE projection of static input embeddings for the 64 standard codons (perplexity = 10), annotated by encoded amino-acid identity. **B**, t-SNE projection of static codon embeddings annotated by amino-acid physicochemical class. **C**, Pairwise cosine similarity between codon embeddings calculated in the original embedding space. Synonymous codon pairs show greater similarity than nonsynonymous pairs (mean *±* s.d., 0.37 *±* 0.13 versus 0.02 *±* 0.06). **D**, t-SNE projection of contextual sequence representations from one-third of the independent held-out species dataset, filtered to no more than 40% protein identity with the pretraining corpus. **E**, Nearest-centroid species classification using the original sequence representations. Centroids were estimated from one-third of the held-out dataset and evaluated on the remaining two-thirds.

At the codon level, the two-dimensional projections revealed clear organization by encoded amino-acid identity and physicochemical class, with synonymous codons tending to occupy neighbouring regions in the embedding space (Fig. 2A,B). Consistent with this organization, pairwise cosine similarity calculated directly in the original embedding space was substantially higher for synonymous than for nonsynonymous codon pairs (mean *±* s.d., 0.37 *±* 0.13 versus 0.02 *±* 0.06; Fig. 2C). Thus, despite being pretrained without explicit amino-acid or biochemical labels, CodonMamba learned codon representations that recapitulate established genetic-code and biochemical relationships.

We next asked whether this organization extended to contextual sequence-level representations. Following the held-out species evaluation strategy introduced by CaLM [32], we analysed an independent dataset whose sequences shared no more than 40% protein identity with the pretraining corpus. The resulting embeddings formed distinct species-associated regions, with partial overlap between the two yeasts, *Saccharomyces cerevisiae* and *Pichia pastoris* (Fig. 2D). To quantify this information, we trained a nearest-centroid classifier on embeddings from one-third of the held-out dataset and evaluated it on the remaining two-thirds. CodonMamba achieved the highest observed classification accuracy among the evaluated models (91.4%), compared with 90.0% for CaLM and 86.1% for mRNAFM [33] (Fig. 2E).

Together, these analyses demonstrate that CodonMamba learns structured coding-sequence representations capturing synonymous codon relationships, amino-acid biochemical organization and species-associated sequence signatures.These biologically informative representations provide a foundation for both downstream mRNA function prediction and CDS-design framework established by CodonMamba.

### 2.2 CodonMamba transfers broadly across diverse mRNA prediction tasks

To test whether the biological structure observed in the pretrained representations translates into functional utility, we evaluated CodonMamba across 12 public mRNA prediction benchmarks spanning protein expression, mRNA stability and degradation, and regulatory functions (Supplementary Table S1). The benchmark panel includes heterogeneous sequence sources and experimental settings, enabling evaluation of whether a shared coding-sequence representation can generalize across distinct mRNA functions. CodonMamba was compared with 11 baseline models spanning codon- and nucleotide-level mRNA models, DNA and RNA foundation models, protein language models, and one-hot sequence representations (Supplementary Table S2).

Under a standardized downstream evaluation protocol, CodonMamba ranked first on 10 of the 12 tasks and second on the remaining two (Fig. 3B; Supplementary Table S3), making it the only evaluated model to rank within the top two across the complete benchmark panel. This cross-task consistency indicates that the pretrained representation captures transferable coding-sequence features underlying diverse mRNA functions rather than task-specific sequence–function associations.

**Figure 3.**
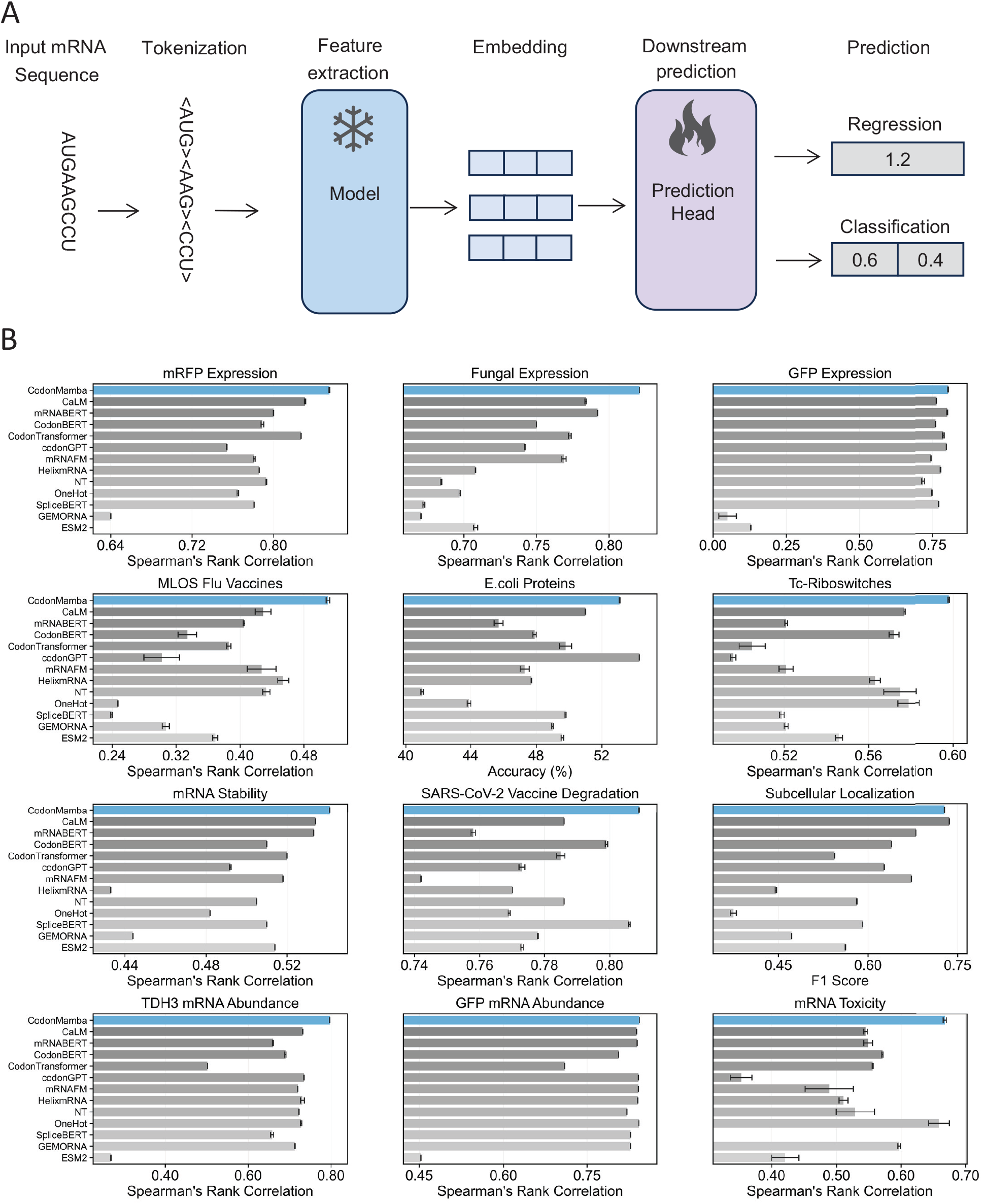
CodonMamba shows broad and consistent transfer across diverse mRNA prediction tasks. **A**, Downstream evaluation framework using frozen pretrained representations and lightweight task-specific prediction heads. **B**, Performance across 12 mRNA prediction benchmarks spanning expression, stability, degradation and regulatory functions. CodonMamba was compared with 11 pretrained baselines spanning codon- and nucleotide-level mRNA models, DNA and RNA foundation models, protein language models, together with one-hot sequence representations. Bars show the mean across three independent runs and error bars denote s.e.m. Undefined and negative Spearman correlations were displayed as zero for visualization only; the underlying values are reported in Supplementary Table S3.

The breadth of this performance extended across model modalities. CodonMamba consistently outperformed models pretrained primarily on DNA, non-coding RNA or protein sequences, supporting the value of coding-sequence-specific pretraining for mRNA prediction. In particular, representations derived from protein language models showed substantially weaker performance on several codon-sensitive tasks, consistent with the loss of synonymous information when coding sequences are reduced to amino-acid sequences. These results support the premise that codon-level representations retain regulatory and translational information that is inaccessible to amino-acid-only models.

CodonMamba also showed more consistent transfer than existing codon language models. codonGPT [25] achieved the highest score on the *E. coli* Proteins benchmark and CaLM led on Subcellular Localization, whereas CodonMamba ranked second on both and achieved the best performance on the remaining ten tasks. Other models showed stronger task-to-task variation, with high performance in selected settings but larger losses elsewhere. By contrast, CodonMamba remained close to the task-specific optimum throughout the benchmark suite.

Together, these results demonstrate that self-supervised codon-level pretraining enables a broadly transferable coding-sequence representation that supports mRNA function prediction rather than specialization to a limited prediction setting. This predictive capability, together with the learned coding-sequence representation, establishes the foundation for CodonMamba’s programmable framework for CDS design.

### 2.3 CodonMamba enables programmable CDS design through inference-time steering

To translate the pretrained coding-sequence distribution into context-configurable CDS design, we formulated generation as constrained synonymous decoding with an inference-time host preference (Fig. 4A). The framework separates the learned coding-sequence distribution from configurable design preferences through two complementary mechanisms. Synonymous-codon masking acts as a hard constraint that restricts each decoding step to codons encoding the target amino acid, thereby defining the protein-preserving design space. Within this admissible space, a host-specific codon usage prior acts as a soft preference that shifts the relative likelihood of synonymous codons without modifying the pretrained model parameters.

**Figure 4.**
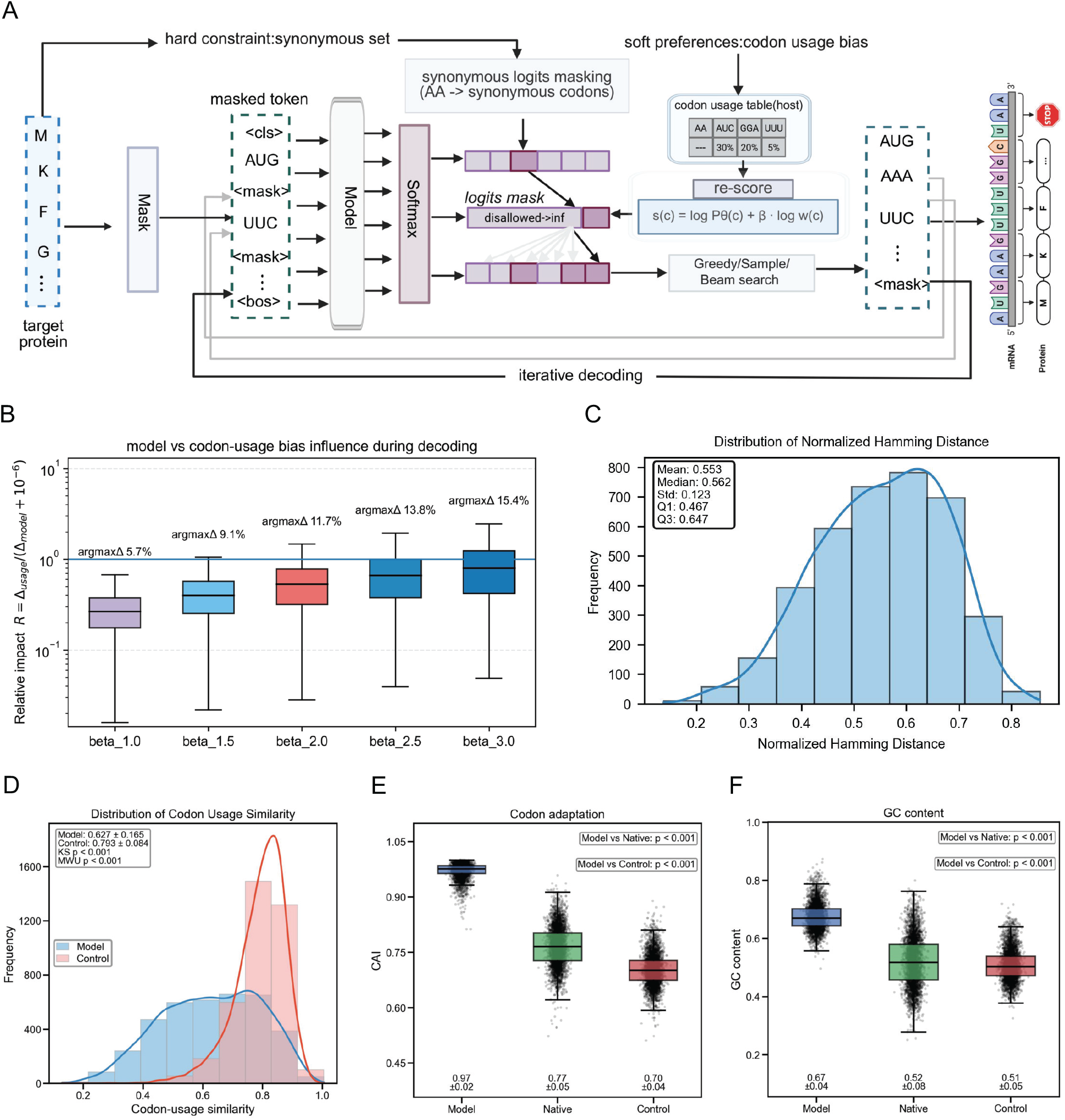
CodonMamba enables programmable CDS design through inference-time steering. **A**, Constrained synonymous decoding by iterative masked codon infilling. Synonym-aware logit masking restricts each position to codons encoding the target amino acid, whereas a host codon usage prior provides a soft preference among the admissible codons. **B**, Influence of prior strength (*β* = 1.0–3.0). *R* is the ratio of the within-synonym logit range introduced by the host prior to that supplied by the pretrained model, and argmaxΔ is the fraction of decoded positions for which introduction of the prior changes the greedy codon choice. **C**, Normalized codon Hamming distance between natural and CodonMamba-generated CDSs at *β* = 2.0. **D**, Native codon-usage similarity of CodonMamba-generated CDSs and substitution-load-matched random synonymous controls. **E**, Human codon adaptation index (CAI) of CodonMamba-generated, natural and matched-random CDSs. **F**, GC content of the same sequence groups.

We evaluated this framework on 3,765 proteins using natural CDSs from Zhang et al. [26] as references and generated synonymous CDSs by greedy decoding with a human codon usage prior. As expected from the hard synonymous constraint, all 3,765 generated sequences preserved the target amino acid sequence, confirming implementation-level fidelity across the complete benchmark set.

We next quantified how strongly the explicit prior perturbs the pretrained model distribution. We varied the prior strength from *β* = 1.0 to 3.0 and measured both the relative strength of the prior within each synonymous codon set (*R*) and the fraction of positions at which introducing the prior changed the greedy codon choice (argmaxΔ; Fig. 4B). Across all tested settings, the prior-derived within-synonym logit range was generally smaller than the corresponding model-derived range (median *R <* 1), whereas the prior changed only 5.7%–15.4% of codon decisions. At *β* = 2.0, the prior altered 11.7% of positions, providing a measurable host-directed effect while leaving most model-preferred codon choices unchanged. We therefore used *β* = 2.0 for subsequent analyses.

Despite the limited effect of the host prior on individual decoding decisions, CodonMamba-generated CDSs diverged substantially from their native references. At *β* = 2.0, the mean normalized codon Hamming distance was 0.553 (median, 0.562; Fig. 4C), corresponding to synonymous differences at more than half of codon positions. Notably, substantial native-referenced divergence was already present under host-free decoding (*β* = 0; Supplementary Fig. S2), whereas introducing the human codon usage prior at *β* = 2.0 changed only 11.7% of model-preferred codon choices. These results indicate that extensive synonymous remodeling is driven primarily by the pretrained CodonMamba distribution, with the host prior providing a comparatively sparse directional adjustment.

To determine whether this recoding reflected systematic codon selection rather than synonymous substitution burden alone, we constructed random synonymous controls matched to each CodonMamba design for the number of substitutions relative to the natural CDS. Despite the matched substitution load, CodonMamba-generated CDSs showed lower similarity to native gene-level codon usage than the random controls (Fig. 4D), indicating that the synonymous changes introduced by the model were non-random.

The resulting sequence shift was strongly aligned with the human codon-usage reference. CodonMamba-generated CDSs achieved a mean human codon adaptation index (CAI) of 0.97, compared with 0.77 for natural CDSs and 0.70 for matched random controls (Fig. 4E). Mean GC content increased in parallel (0.67, 0.52 and 0.51, respectively; Fig. 4F), showing that host-directed decoding altered both synonymous codon usage and broader nucleotide composition. Consistently, comparison of host-free (*β* = 0) and human-prior-guided (*β* = 2.0) decoding showed a pronounced increase in human CAI but only modest changes in native-referenced divergence (Supplementary Fig. S2).

Together, these results distinguish extensive model-driven synonymous remodeling from the more limited directional effect of the host codon usage prior. Synonym-aware masking preserves the encoded protein, whereas the replaceable prior redirects the pretrained sequence distribution towards the specified host preference at inference.

### 2.4 CodonMamba enables coordinated multi-property CDS design

The preceding analyses distinguished extensive model-driven synonymous remodeling from the more limited directional effect of the host codon usage prior. We next examined whether CodonMamba-generated CDSs occupy sequence regimes associated with experimentally measured mRNA properties, how these properties respond to inference-time host steering, and how the resulting design profiles compare with existing CDS design approaches.

We first assessed a CodonMamba-derived naturalness score, defined from the length-normalized pseudo-log-likelihood assigned by the pretrained masked language model. Higher naturalness therefore reflects greater compatibility with coding-sequence contexts assigned high likelihood by CodonMamba, following likelihood-based sequence scoring used previously in biological language models and mRNA design [26, 34, 35]. We evaluated this score against experimentally measured protein expression and mRNA stability in three independent datasets compiled by Zhang et al. [26], spanning distinct reporter systems, experimental settings and mRNA modification states (Fig. 5A; see Methods). None of these datasets was used during CodonMamba pretraining or for task-specific fitting in this analysis.

**Figure 5.**
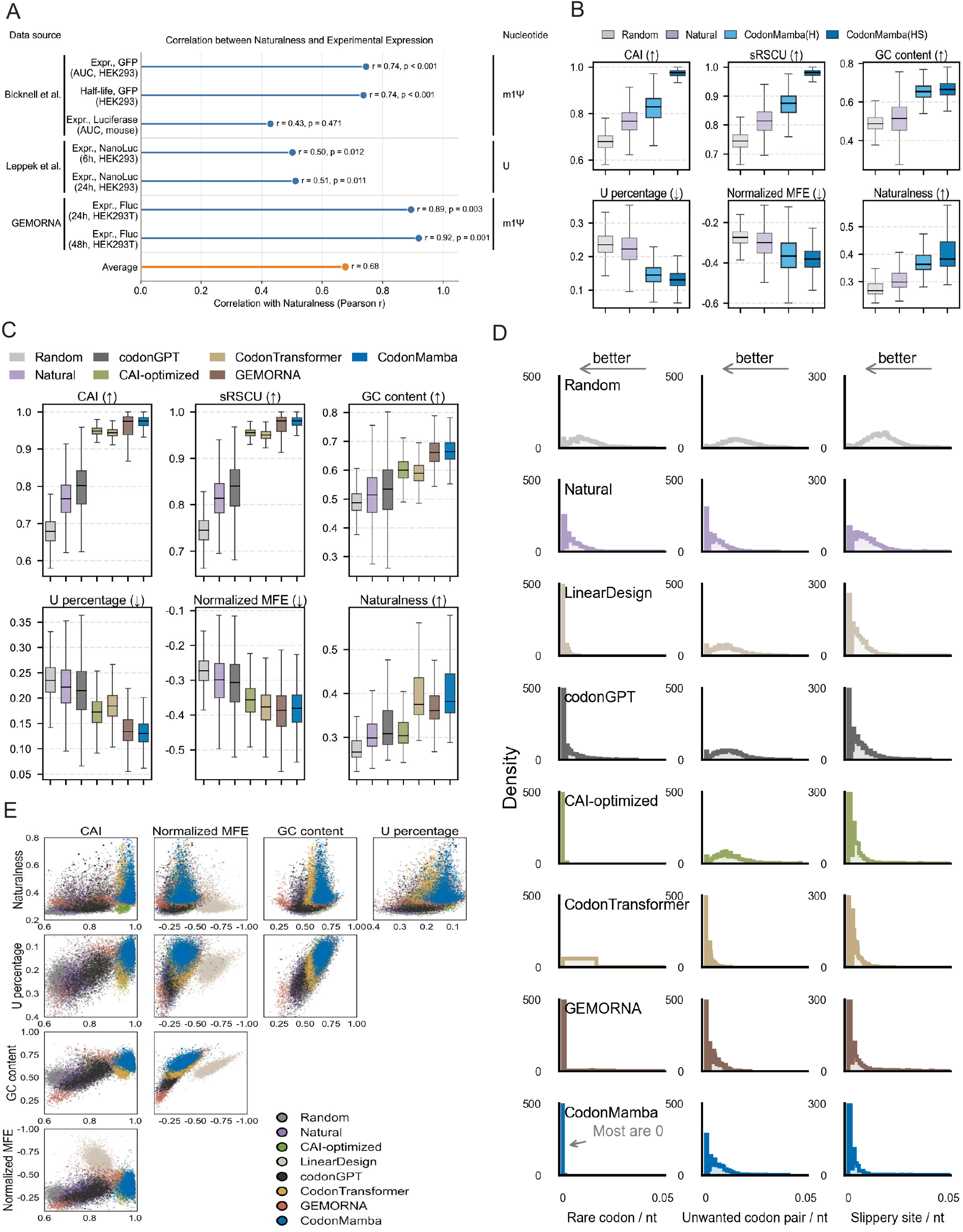
CodonMamba enables coordinated multi-property CDS design. **A**, Pearson correlations between CodonMamba-derived naturalness and experimentally measured mRNA expression or stability across independent datasets. Naturalness was calculated from the CodonMamba model. **B**, Six sequence properties for host-free decoding (H, *β* = 0) and decoding with a human codon usage prior (HS, *β* = 2.0). Comparisons were paired by target protein and assessed using two-sided Wilcoxon signed-rank tests. **C**, Comparison of six global sequence properties across CodonMamba and representative CDS design baselines. **D**, Rare-codon, unwanted codon-pair and slippery-site densities across sequence groups. **E**, Pairwise feature-space distributions for representative CDS design methods.

Naturalness was positively associated with the experimental measurements across the evaluated datasets, with a mean Pearson correlation of *r* = 0.68 and correlations exceeding *r* = 0.9 for the strongest readouts (Fig. 5A). Its overall association with experimental measurements was comparable in magnitude to that of sequence-level properties including GC and uridine content (Supplementary Fig. S3). Notably, CodonMamba-generated sequences showed higher model-derived naturalness than conventional optimization approaches, suggesting that the inferred redesigns remained compatible with the broader coding-sequence distribution learned during pretraining.

We next examined how inference-time host steering reshaped broader sequence properties. Using the same 3,765 target proteins, we compared host-free decoding (H, *β* = 0) with decoding guided by a human codon usage prior (HS, *β* = 2.0) across six properties spanning codon adaptation, nucleotide composition, predicted RNA structure and model-derived naturalness (Fig. 5B). These properties also showed measurable associations with experimental expression and stability in the external datasets (Supplementary Fig. S3), supporting their use as complementary design readouts rather than direct measures of functional performance.

Across *β* = 0–3.0, increasing prior strength progressively increased CAI and sRSCU, reduced uridine content whereas normalized MFE showed a comparatively smaller change in magnitude (Supplementary Fig. S4). Most of these changes occurred between *β* = 0 and *β* = 2.0, with comparatively smaller additional shifts at higher prior strengths. Naturalness remained broadly stable across the same range, indicating that substantial changes in codon adaptation and nucleotide composition could be introduced without a correspondingly large change in compatibility with the pretrained model distribution. At *β* = 2.0, median CAI increased from 0.828 to 0.976, a 17.9% relative increase, and the other evaluated properties also differed significantly between paired designs for the same proteins (two-sided paired Wilcoxon signed-rank tests, all *P <* 0.001). These property-level changes accompanied alterations at only 11.7% of model-preferred codon choices (Fig. 4B), showing that a relatively limited set of prior-induced decoding changes can propagate to broader sequence-level properties.

We then compared CodonMamba with natural CDSs, random synonymous controls, classical CAI optimization and LinearDesign[16], and the generative models codonGPT[25], CodonTransformer[36] and GEMORNA [26]. Following previous CDS design evaluations [26], we considered complementary properties spanning host codon adaptation, nucleotide composition, predicted RNA secondary structure, model-derived naturalness and local codon-context features (Fig. 5C–E). The corresponding six-property profile for LinearDesign is shown separately in Supplementary Fig. S5.

Across the methods shown in Fig. 5C,CodonMamba achieved host codon adaptation comparable to specialized optimization methods while maintaining higher compatibility with its learned coding-sequence distribution. Both measures were higher than those observed for codonGPT, CAI-optimized and CodonTransformer sequences. CodonMamba and GEMORNA also showed relatively high GC content and low uridine content, whereas natural, random and codonGPT sequences occupied lower-GC and higher-uridine ranges. LinearDesign showed lower CAI and sRSCU values than CodonMamba (Supplementary Fig. S5), reflecting its joint optimization of codon usage and RNA secondary-structure free energy rather than optimization of codon adaptation alone.

The methods also differed in predicted RNA structural properties. Unlike LinearDesign, which explicitly optimizes RNA secondary structure free energy and produced the most negative MFE distribution, CodonMamba generated sequences within a structural regime comparable to GEMORNA and CodonTransformer (Fig. 5C). LinearDesign, by contrast, produced a substantially more negative normalized MFE distribution (Supplementary Fig. S5), consistent with its explicit optimization of RNA secondary-structure free energy [16]. Thus, CodonMamba altered predicted structural properties without the pronounced MFE shift produced by LinearDesign. CodonMamba also showed naturalness values among the highest of the evaluated methods. Because this score is derived from CodonMamba itself, however, this comparison reflects compatibility with the learned coding-sequence distribution rather than independent evidence of functional superiority.

Differences extended to local codon context (Fig. 5D). In contrast to CAI-driven optimization, which increased unwanted codon-pair density, CodonMamba maintained codon-context features closer to natural sequences. CodonMamba designs also showed low rare-codon and slippery-site densities, indicating that strong host codon adaptation was not accompanied by marked accumulation of these local sequence features.

Finally, pairwise feature-space analyses revealed distinct multi-property profiles across design strategies (Fig. 5E; Supplementary Fig. S6). Classical optimization methods showed more objective-specific shifts: CAI optimization primarily displaced sequences towards high codon adaptation, whereas LinearDesign occupied a distinct region associated with stronger RNA-structure optimization. Generative approaches occupied broader multi-property regions, with CodonMamba combining strong host codon adaptation with coordinated nucleotide composition, predicted RNA structure and local codon-context properties. Together, these results demonstrate that CodonMamba performs coordinated multi-property CDS redesign by combining host-adaptive codon usage with preservation of learned sequence characteristics, rather than optimizing a single predefined objective.

### 2.5 CodonMamba enables programmable cross-host CDS redesign through inference-time steering

We next asked whether a single pretrained CodonMamba model could be redirected across expression hosts by replacing only the inference-time codon usage prior. Using the same 3,765 target proteins, we generated synonymous CDSs with Human, Mouse, *E. coli* and Yeast codon usage priors at *β* = 2.0, while keeping the pretrained model parameters unchanged.

Across all four host conditions, the prior-derived within-synonym logit range was generally smaller than the corresponding model-derived range (median *R <* 1), and introducing the prior altered only 11.2%–13.9% of model-preferred codon choices (Fig. 6A). The *E. coli* and Yeast priors produced somewhat larger changes than the Mouse prior, but most decoding decisions remained unchanged in every condition. Thus, switching the host prior redirected codon selection through a relatively limited subset of synonymous decisions.

**Figure 6.**
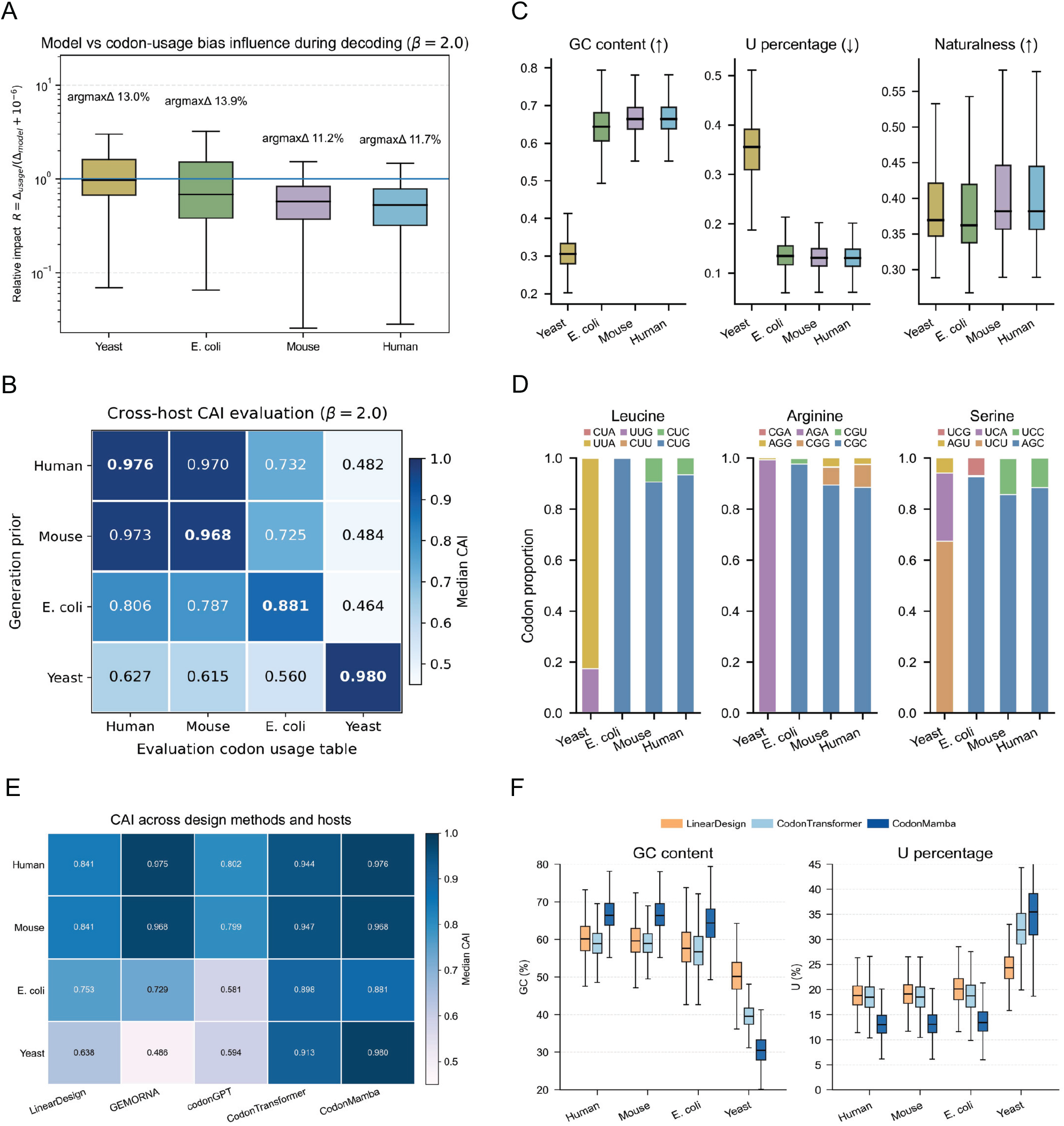
CodonMamba enables programmable cross-host CDS redesign through inference-time steering. **A**, Relative influence of the pretrained model and host codon usage prior during decoding with Human, Mouse, *E. coli* and Yeast priors (*β* = 2.0). *R* denotes the prior-to-model within-synonym logit-range ratio; percentages indicate argmaxΔ. **B**, Cross-host codon adaptation index (CAI). Rows indicate the host prior used for generation and columns the host codon usage reference used for evaluation; values denote median CAI. **C**, GC content, uridine percentage and CodonMamba-derived naturalness across the four host-prior conditions. **D**, Synonymous codon usage for the six-fold degenerate amino acids leucine, arginine and serine across host-prior conditions. **E**, Host-matched median CAI across representative computational CDS design methods. GEMORNA and codonGPT were treated as host-independent outputs in this comparison and evaluated against each host-specific reference without regeneration. **F**, GC content and uridine percentage under matched-host settings for methods with host- or species-dependent CDS designs.

Despite this limited intervention, codon usage was systematically redirected towards the corresponding host. Cross-host CAI showed a pronounced diagonal-dominant pattern, with matched-host median values of 0.976 for Human, 0.968 for Mouse, 0.881 for *E. coli* and 0.980 for Yeast (Fig. 6B). Human- and Mouse-prior designs remained similar under reciprocal evaluation, whereas stronger separation was observed for the more divergent *E. coli* and Yeast references. Relative to host-free decoding, the *E. coli* prior increased *E. coli*-reference CAI from 0.672 to 0.881 (31%), whereas the Yeast prior increased Yeast-reference CAI from 0.463 to 0.980 (112%). These shifts were associated with changes at only 13.9% and 13.0% of model-preferred codon choices, respectively. Codon-usage cosine similarity showed the same diagonal-dominant organization (Supplementary Fig. S7), indicating that prior switching reshaped the broader synonymous codon distribution rather than only increasing a scalar adaptation metric.

Host-dependent changes extended to global nucleotide composition (Fig. 6C). Human- and Mouse-prior designs were relatively GC-rich and U-poor, whereas the Yeast prior shifted the same protein designs towards lower GC and higher uridine content, consistent with the corresponding species-level codon preferences. CodonMamba-derived naturalness was highest under the Human and Mouse priors and lower under the *E. coli* and Yeast conditions, indicating greater departure from sequence patterns assigned high likelihood by the pretrained model under the more divergent host priors.

The same host dependence was evident within individual synonymous codon families (Fig. 6D). For leucine, Yeast-prior designs preferentially used UUG, whereas *E. coli*-prior designs strongly favoured CUG; Human- and Mouse-prior designs also favoured CUG but retained a broader codon distribution. For arginine, *E. coli*-prior designs preferentially used CGC and CGU, whereas Yeast-prior designs favoured AGA. These amino-acid-specific shifts were consistent with the corresponding changes in CAI, codon-usage profiles and nucleotide composition.

We next compared host-matched CAI across representative CDS design methods (Fig. 6E). LinearDesign was evaluated under corresponding host-specific optimization settings, whereas CodonTransformer used species-conditioned generation. GEMORNA and codonGPT were treated as host-independent outputs for a given target protein and were therefore evaluated against each host reference without regeneration. CodonMamba achieved high or comparable matched-host CAI across all four host references using the same pretrained model with only the external codon usage prior replaced. Host-dependent nucleotide composition was similarly evident in comparisons with LinearDesign and CodonTransformer (Fig. 6F): CodonMamba produced relatively GC-rich, U-poor sequences under the Human, Mouse and *E. coli* priors, whereas the Yeast prior shifted the same protein designs towards lower GC and higher uridine content.

CAI-optimized sequences provided an additional positive control and achieved high matched-host CAI by construction (Supplementary Fig. S8). Because CAI was both an explicit design criterion and the evaluation metric, these sequences were not treated as an independent generative benchmark.

Together, these results show that replacing a single inference-time codon usage prior is sufficient to redirect synonymous codon usage and broader sequence composition across distinct host references without retraining the CodonMamba backbone.

## 3 Discussion

CodonMamba connects general coding-sequence representation learning with constrained and programmable CDS design. Its central principle is to retain the contextual sequence regularities learned during pretraining while enabling application-specific preferences to directly steer generation at inference. Large-scale self-supervised pretraining provides a contextual distribution over natural coding sequences, synonymous-codon masking restricts generation to a protein-preserving design space, and a replaceable codon usage prior adjusts the relative preference among admissible synonymous codons without modifying the pretrained model parameters. Together, these components allow generation to be redirected towards an intended application context while retaining both the encoded protein and the sequence information captured by the pretrained backbone.

The representation and downstream analyses support codon-level pretraining as a biologically informative basis for this framework. CodonMamba recapitulated established relationships among synonymous codons, amino-acid physicochemical classes and species-associated coding-sequence patterns, and transferred consistently across 12 heterogeneous mRNA prediction benchmarks. These results do not imply that codon language modelling newly reveals the organization of the genetic code; rather, they show that a bidirectional state-space model retains biologically meaningful information previously observed in codon-based representations while supporting broad transfer across distinct mRNA-related functions. Its performance relative to models pretrained on DNA, non-coding RNA and protein sequences further supports direct modelling of CDSs, where synonymous variation preserves regulatory and translational information that is lost when coding sequences are reduced to amino-acid sequences.

The generation experiments show how direct inference-time steering interacts with the coding-sequence distribution learned during pretraining. Substantial synonymous remodeling was already produced under host-free decoding, whereas introducing a host codon usage prior changed only a minority of the codon choices preferred by the pretrained model. The prior therefore does not replace the learned sequence distribution with a conventional codon-optimization rule, but selectively redirects synonymous decisions within that distribution. Despite this limited intervention, replacing only the external prior systematically retargeted the same pretrained model towards human, mouse, *E. coli* and yeast codon preferences, with corresponding changes in codon usage and nucleotide composition. The coordinated changes observed in codon adaptation, predicted RNA structure and local codon context further suggest that the pretrained model captures dependencies extending beyond any single explicitly specified design objective. Programmability here therefore refers not simply to generating alternative synonymous CDSs, but to systematically reconfiguring the sequence preference applied to the same learned coding-sequence distribution at inference.

This formulation complements rather than replaces existing CDS design paradigms. Classical approaches such as CAI optimization provide precise and interpretable control over predefined codon-usage objectives, whereas LinearDesign explicitly incorporates RNA secondary-structure stability into sequence optimization [**?**]. Generative approaches can capture broader higher-order dependencies from natural sequence data, but application-specific control is commonly introduced through training-time conditioning, model optimization or additional guidance mechanisms. Recent protein-generation methods have similarly demonstrated generation-time control through activation-based steering or predictor-based property guidance [29, 30]. CodonMamba implements a distinct form of inference-time control: an externally specified codon usage prior acts directly on the relative logits of admissible synonymous codons within an explicitly protein-preserving design space. This allows application-specific preferences to be replaced without retraining the generative backbone or requiring an auxiliary property predictor. Host codon usage provides a simple and interpretable demonstration of this strategy, while the underlying formulation permits the generative behaviour of a shared pretrained CDS model to be reconfigured through an external inference-time preference.

A major limitation of the present study is the absence of direct experimental validation. The current analyses therefore establish computational control over protein-preserving sequence generation, codon-usage retargeting and associated sequence properties, but do not demonstrate that CodonMamba-designed CDSs improve protein expression, mRNA stability or other functional outcomes in specific biological systems. Direct experimental comparison of generated CDSs will be important for determining how these computational design properties translate into biological performance.

Several directions could extend the current framework. Future inference-time priors could incorporate more biologically resolved signals associated with particular tissues, cell types or cell lines, including context-specific codon-usage patterns, tRNA availability or experimentally measured translation efficiency. Such signals could, in principle, redirect the same pretrained CodonMamba backbone towards distinct translational environments without repeated model retraining, although their value as design priors would require independent experimental calibration. More complex sequence preferences, including RNA structural properties or co-translational-folding-related constraints, could also be incorporated into the decoding process. Extending the framework to multiple simultaneously acting preferences would require careful characterization of interactions and trade-offs among the corresponding control signals.

More broadly, CodonMamba illustrates a strategy for biological sequence generation in which the contextual information captured by large-scale pretraining is retained while application-specific preferences are introduced directly during generation. Here, this principle is demonstrated through protein-preserving synonymous CDS generation and programmable cross-host codon-usage retargeting. With direct experimental validation and more biologically resolved control signals, this framework could provide a flexible route towards context-configurable CDS design without repeated model retraining.

## 4 Methods

### Datasets

#### Pretraining corpus

We used the processed coding-sequence corpus released with CaLM [32] for CodonMamba pretraining. The final corpus contains 8,771,938 coding sequences derived from genomes deposited in the European Nucleotide Archive (ENA; data snapshot, April 2022) [37] and spans 1,544 organisms across the tree of life. In the original CaLM preprocessing pipeline, sequences containing ambiguous nucleotides, non-AUG start codons, internal stop codons or lengths not divisible by three were excluded. Sequences were subsequently grouped by organism, translated to amino-acid sequences and clustered at 40% amino-acid identity using CD-HIT [38] to reduce redundancy while retaining phylogenetic diversity [38]. We used the resulting processed corpus without reconstructing the original ENA collection. Sequences exceeding the maximum CodonMamba context length were truncated to 1,024 codon tokens before pretraining.

#### Downstream prediction datasets

We evaluated pretrained representations on 12 public mRNA prediction datasets spanning protein expression (five datasets), mRNA stability and degradation (four datasets), and regulatory phenotypes (three datasets). Processed benchmarks were drawn primarily from the collections released with CodonBERT [24], CaLM [32] and SynCodonLM [39], which curate data from previous experimental studies and public resources. Collectively, the datasets comprise endogenous genes and exogenous reporter constructs from multiple species and experimental settings, including both unmodified and m^1^Ψ-modified mRNAs, and range from 167 to 65,356 sequences.

Author-provided training, validation and test partitions were retained where available to preserve comparability with previous studies. For datasets without predefined partitions, sequences were divided into training, validation and test sets at a 0.70/0.15/0.15 ratio. To ensure consistent input lengths across models, sequences exceeding 1,024 codons were truncated to 1,024 codons before representation extraction. Original data sources, dataset-specific provenance, endpoints, sequence lengths and task definitions are summarized in Supplementary Table S1, with additional details on dataset processing, partitioning and evaluation provided in Supplementary Note.

#### Held-out species dataset

Representation diagnostics were performed using the independent held-out dataset released with CaLM [32]. The dataset contains 4,358 coding sequences from seven model organisms: three multicellular eukaryotes (*Arabidopsis thaliana, Drosophila melanogaster* and *Homo sapiens*), two unicellular eukaryotes (*Saccharomyces cerevisiae* and *Pichia pastoris*), one bacterium (*Escherichia coli*) and one archaeon (*Haloferax volcanii*). Construction and homology filtering of this dataset followed the original CaLM protocol, including sequence clustering and removal of homologous sequences from the pretraining corpus [32].

#### Experimental datasets for naturalness evaluation

To evaluate whether model-derived naturalness was associated with experimentally measured mRNA properties, we used three independent datasets compiled by Zhang et al. [26]. The Bicknell dataset contains exogenous GFP mRNAs incorporating m^1^Ψ and measurements obtained in human and mouse systems [40]. The Leppek dataset contains unmodified exogenous mRNAs encoding multiple reporter proteins in human cells [41]. The GEMORNA experimental dataset contains m^1^Ψ-modified mRNAs encoding destabilized firefly luciferase (Fluc2P) evaluated in HEK293T cells [26]. None of these measurements was used during CodonMamba pretraining or for task-specific fitting in the naturalness analysis.

#### Generation benchmark dataset and controls

CDS generation was evaluated on 3,765 mammalian target proteins from the GEMORNA benchmark [26]. natural CDSs, random synonymous controls, GEMORNA-generated sequences, and the human-reference CAI-optimized and LinearDesign sequences were obtained directly from the released benchmark and analysed without regeneration. Additional baseline generation procedures are described in Supplementary Note.

For the substitution-load-controlled analysis in Fig. 4, we constructed an additional matched random synonymous control separately for each target. Each matched control contained the same number of synonymous substitutions relative to the natural CDS as the corresponding CodonMamba design while preserving the encoded amino-acid sequence. This control was used to distinguish systematic model-driven codon selection from effects attributable solely to synonymous substitution burden.

### Model architecture and pretraining

#### Codon tokenization and model architecture

CodonMamba is a bidirectional masked language model operating on non-overlapping codon tokens. Coding sequences were tokenized into triplets using a vocabulary containing the 64 standard codons and five special tokens (`<cls>`, `<eos>`, `<pad>`, `<unk>` and `<mask>`). Tokens were mapped to learned 768-dimensional embeddings.

The backbone comprises eight stacked bidirectional Mamba (BiMamba) blocks with hidden dimension *d* = 768. The model contains approximately 71 million trainable parameters and supports sequences of up to 1,024 codon tokens. An MLM output projection maps the final hidden representations to logits over the codon vocabulary at each sequence position.

#### Bidirectional Mamba block

To incorporate information from both sequence directions, we used the parameter-efficient bidirectional Mamba formulation introduced in Caduceus [42]. Given an input representation *x* ∈ ℝ^*T×d*^, the forward branch is

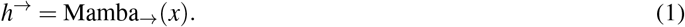

The reverse branch operates on the sequence in reverse order:

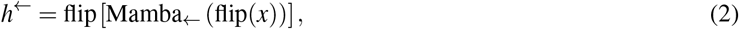

where flip(·) reverses the sequence dimension. The two directional representations are combined by element-wise summation:

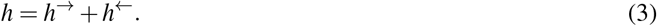

Following the BiMamba implementation used here, projection layers are shared between the two directional branches, whereas the convolutional and state-space components remain direction specific. This reduces the additional parameter cost of bidirectional processing relative to two completely independent Mamba blocks [31, 42].

#### Masked language modelling objective

CodonMamba was pretrained using masked language modelling. For an input sequence **x** = (*x*_1_, …, *x*_*L*_), a subset of positions ℳ was selected and corrupted to produce 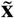. The training objective was

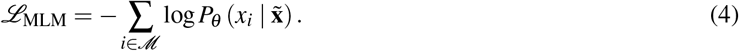

The loss was evaluated only at the selected masking positions.

#### Pretraining procedure

For each training sequence, 15% of codon positions were selected for corruption. Of these selected positions, 75% were replaced by `<mask>`, 15% by a randomly selected vocabulary token and 10% were left unchanged. Sequences within each batch were padded to the length of the longest sequence after truncation to the maximum context length.

Optimization used AdamW [43] with a peak learning rate of 4 *×* 10^−4^, weight decay of 0.01 and *β* parameters (0.9, 0.999). The learning rate was increased linearly during the first 1,500 optimization steps and subsequently decayed using a cosine schedule. Gradients were accumulated over four mini-batches. One per cent of the pretraining corpus was reserved for validation. Training was stopped after 47 epochs when validation loss had plateaued, corresponding to approximately seven days of training on four NVIDIA A100 GPUs.

### Representation and downstream evaluation

#### Codon embedding analysis

Following representation analyses used in previous codon language models [24, 32], static input embeddings for the 64 standard codon tokens were extracted directly from the learned embedding matrix. Special-token embeddings were excluded. For visualization, the 768-dimensional codon embeddings were projected into two dimensions using t-distributed stochastic neighbour embedding (t-SNE; scikit-learn v1.3.2) with `n_components=2` and `perplexity=10`. The same projection was annotated by encoded amino-acid identity and amino-acid physicochemical class.

Codon similarity was quantified independently of the two-dimensional projection by calculating pairwise cosine similarities directly in the original embedding space. Codon pairs encoding the same amino acid were classified as synonymous and compared with nonsynonymous codon pairs.

#### Sequence embedding visualization and species classification

For each sequence in the CaLM held-out dataset, a sequence-level representation was obtained from the final hidden state corresponding to the `<cls>` token. For visualization, sequence representations were projected into two dimensions using t-SNE [44] (scikit-learn v1.3.2) with `n_components=2` and `perplexity=30`; points were annotated according to species.

Species discrimination followed the nearest-centroid protocol introduced by CaLM [32]. The held-out dataset was divided into a centroid-estimation subset containing approximately one-third of the sequences and an independent test subset containing the remaining two-thirds. For each species, the centroid was calculated as the mean sequence representation in the centroid-estimation subset. Each test representation was assigned to the species centroid with the smallest Euclidean distance. Classification accuracy was calculated as the proportion of correctly assigned test sequences.

#### Downstream prediction benchmarks

CodonMamba was evaluated across 12 downstream mRNA prediction benchmarks against representative models spanning codon- and nucleotide-level mRNA models, DNA and RNA foundation models, and protein language models, together with one-hot sequence representations (Supplementary Table S2). For all pretrained models, backbone parameters were frozen and only a lightweight task-specific prediction head was optimized (Fig. 3A). Contextual sequence representations were processed using a shared ResNet-style convolutional head [45] (Supplementary Fig. S1), with the output layer adapted to regression, multi-class classification or multi-label classification according to the benchmark endpoint.

To ensure comparability across models, the same dataset partitions, input-length controls and downstream evaluation protocol were used for each benchmark, with the same prediction-head architecture applied wherever compatible with the representation format. Each model–benchmark combination was evaluated in three independent runs with different random seeds. Reported performance corresponds to the mean across runs, and error bars in Fig. 3 denote s.e.m. Dataset processing, partitioning and evaluation metrics are detailed in Supplementary Note.

### Constrained CDS generation

CodonMamba generates protein-preserving coding sequences through iterative masked infilling. The decoding strategy builds on iterative conditional generation for bidirectional masked language models [46] and adapts multi-position masked decoding, as used in ESM3 [18], to codon-level generation. Protein identity is enforced using the inference-time synonymous logit-masking principle introduced in codonGPT [25], adapted here to parallel masked infilling. A separate host codon usage prior provides optional soft steering at inference. Thus, biological validity and context-specific sequence preference are introduced as distinct decoding components.

Unless otherwise stated, all primary generation experiments in Figs. 4–6 used greedy decoding with a 12-step linear Mask-and-Predict schedule. Alternative stochastic and beam-search decoding strategies are described in Supplementary Note and evaluated in Supplementary Fig. S9.

#### Iterative masked infilling

For a target protein **a** = (*a*_1_, …, *a*_*L*_), generation was initialized from an all-`<mask>` codon sequence of length *L* and proceeded using a 12-step linear Mask-and-Predict schedule. At each iteration, CodonMamba predicted all remaining masked positions, and confidence was defined as the highest probability among the admissible synonymous codons at each position. The highest-confidence positions scheduled for that iteration were decoded by greedy selection and fixed for subsequent iterations. Decoding continued until all positions had been assigned.

#### Hard constraint: synonym-aware logit masking

Protein identity was enforced by adapting the inference-time synonymous logit-masking strategy introduced in codonGPT [25] to iterative masked infilling. For each target amino acid *a*_*i*_, decoding was restricted to its synonymous codon set *V* (*a*_*i*_) using

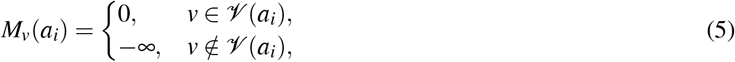

such that the constrained logits were

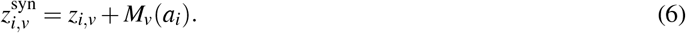

Non-admissible codons therefore have zero probability during decoding, guaranteeing preservation of the target amino-acid sequence.

#### Soft preference: host codon usage prior

Host-specific codon preferences were introduced at inference independently of the pretrained model parameters. Species-level codon usage frequencies for *Homo sapiens* (Human), *Mus musculus* (Mouse), *Escherichia coli* and *Saccharomyces cerevisiae* (Yeast) were obtained from the Codon Usage Database (Kazusa) [47]. For each codon *c* encoding amino acid *a*, a relative adaptiveness weight was defined within its synonymous codon family as

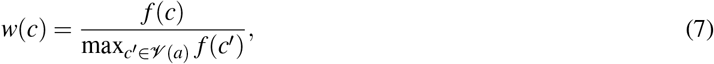

where *f* (*c*) denotes the corresponding species-level codon usage frequency.

Following synonym-aware masking, the logit of each admissible codon *v* ∈ *V* (*a*_*i*_) was adjusted as

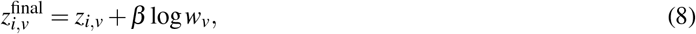

where *z*_*i,v*_ is the CodonMamba logit and *β* ≥ 0 controls the strength of the host preference. Thus, *β* = 0 corresponds to host-free decoding, whereas increasing *β* progressively biases synonymous codon selection towards the supplied host-specific usage profile without modifying the pretrained model parameters.

#### Quantifying inference-time prior strength

To quantify the relative influence of the pretrained model and the host codon usage prior during decoding, we used two complementary measures. For each decoded position, we compared the dynamic range of the model logits with that introduced by the host prior across the admissible synonymous codons:

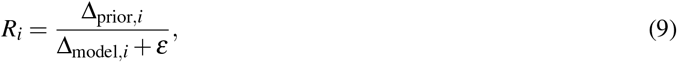

where Δ_model,*i*_ and Δ_prior,*i*_ denote the within-synonym logit ranges contributed by the pretrained model and the host prior, respectively, and *ε* = 10^−6^. Thus, *R*_*i*_ *<* 1 indicates that the prior introduces a smaller within-set logit contrast than the pretrained model.

We additionally quantified the realized effect of the prior as argmaxΔ, defined as the fraction of decoded positions for which adding the host prior changed the greedy synonymous-codon choice relative to host-free decoding:

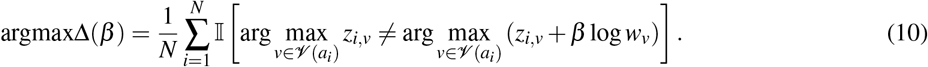

Prior strength was evaluated at *β* ∈ {0, 1.0, 1.5, 2.0, 2.5, 3.0} . Unless otherwise stated, *β* = 2.0 was used for the primary host-guided generation analyses. Detailed definitions and treatment of single-codon amino acids are provided in Supplementary Note; sensitivity analyses are shown in Supplementary Fig. S4.

#### Cross-host prior switching

Programmable host retargeting was evaluated using Human, Mouse, *E. coli* and Yeast codon usage priors. The same pretrained CodonMamba parameters and the same 3,765 target proteins were used for all four conditions; only the external codon usage prior was replaced. Generation used *β* = 2.0 in all matched-host analyses. Each generated sequence set was evaluated against all four host references using CAI and codon-usage profile cosine similarity, enabling matched- and mismatched-host comparisons without model retraining.

### Generated-sequence evaluation

#### CodonMamba naturalness score

Sequence naturalness was quantified using pseudo-log-likelihood (PLL), a standard sequence-scoring procedure for masked language models [48]. Naturalness was calculated using the base pretrained CodonMamba model without task-specific fitting. For a CDS **S** = (*x*_1_, …, *x*_*L*_), each codon was masked individually and scored conditional on the remaining sequence:

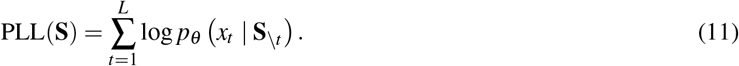

The length-normalized CodonMamba naturalness score was defined as

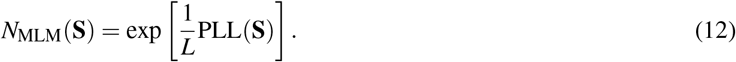

Higher values indicate greater compatibility with the coding-sequence distribution assigned high likelihood by the pretrained CodonMamba model. Naturalness was evaluated against the independent experimental datasets described above using Pearson correlation with the corresponding expression or stability measurements.

#### Sequence-level design metrics

Generated CDSs were evaluated using complementary metrics describing protein fidelity, sequence divergence, host codon matching, nucleotide composition, predicted RNA structure and local codon context. Protein fidelity was assessed by translating each generated CDS and comparing the resulting amino-acid sequence with the target protein.

Divergence from natural CDSs was quantified using normalized codon Hamming distance, synonymous substitution number, codon match rate and gene-level codon-usage similarity. Host matching was assessed using codon adaptation index (CAI) and transcript-level scaled relative synonymous codon usage (sRSCU). Additional sequence properties included GC content, uridine percentage, normalized minimum free energy (nMFE), rare-codon density, unwanted codon-pair density and slippery-site density.

Metrics were calculated at the gene level after sequence cleaning and codon tokenization. Incomplete terminal triplets were removed, paired sequences were restricted to their shared codon-aligned length where required and stop codons were excluded from codon-usage and host-matching calculations. Detailed definitions, normalization procedures and reference sources for all generated-CDS metrics are provided in Supplementary Note.

#### Comparison with CDS design baselines

CodonMamba-generated sequences were compared with natural CDSs, random synonymous controls, CAI-based optimization, LinearDesign and the generative models codonGPT, Codon-Transformer and GEMORNA [16, 25, 26, 36].

For the human-reference comparisons in Fig. 5, natural CDSs, random synonymous controls, GEMORNA-generated sequences, CAI-optimized sequences and LinearDesign designs were obtained directly from the GEMORNA benchmark [26]. codonGPT and CodonTransformer sequences were generated from the same target proteins using the publicly released source code and pretrained models provided by the respective authors.

For the cross-host comparisons in Fig. 6, the human CAI-optimized and LinearDesign sequences were taken directly from the GEMORNA benchmark. Corresponding Mouse, *E. coli* and Yeast CAI-optimized and LinearDesign sequences were generated separately using the codon-usage reference for each target species. CodonTransformer was generated using the corresponding supported species condition. GEMORNA and codonGPT outputs were treated as host-independent sequences for a given target protein in this comparison and were therefore evaluated against different host references without regeneration. Baseline-specific generation procedures are described in Supplementary Note.

## Code and data availability

The CodonMamba model is available at Hugging Face (https://huggingface.co/langmei/CodonMamba). The most updated code and data can be found on GitHub at https://github.com/meilanglang/CodonMamba.

## Acknowledgements

This work was supported in part by the Key Research and Development Program of Zhejiang (2025C01129 to X.L.) and the China Postdoctoral Science Foundation (2025M772773 to Z.W.).

## Author contributions

X.L., K.Y.T. and J.Z. supervised the study, acquired funding, and provided resources. M.L. and X.L. conceptualized the study, developed the methodology, conducted the investigation, curated data, performed formal analysis and visualization, developed the software, and wrote the original draft. All authors reviewed and edited the manuscript.

## Conflict of interest

All authors declare no financial interest.

## Notes

### Competing Interest Statement

The authors have declared no competing interest.

## References

[1] Norbert Pardi, Michael J. Hogan, Frederick W. Porter, and Drew Weissman. mRNA vaccines — a new era in vaccinology. Nature Reviews Drug Discovery, 17(4):261–279, April 2018.

[2] Fernando P. Polack, Stephen J. Thomas, Nicholas Kitchin, Judith Absalon, Alejandra Gurtman, Stephen Lockhart, John L. Perez, Gonzalo Pérez Marc, Edson D. Moreira, Cristiano Zerbini, Ruth Bailey, Kena A. Swanson, Satrajit Roychoudhury, Kenneth Koury, Ping Li, Warren V. Kalina, David Cooper, Robert W. Frenck, Laura L. Hammitt, Özlem Türeci, Haylene Nell, Axel Schaefer, Serhat Ünal, Dina B. Tresnan, Susan Mather, Philip R. Dormitzer, Uğur Ş ahin, Kathrin U. Jansen, and William C. Gruber. Safety and Efficacy of the BNT162b2 mRNA Covid-19 Vaccine. The New England Journal of Medicine, page NEJMoa2034577, December 2020.

[3] Lindsey R. Baden, Hana M. El Sahly, Brandon Essink, Karen Kotloff, Sharon Frey, Rick Novak, David Diemert, Stephen A. Spector, Nadine Rouphael, C. Buddy Creech, John McGettigan, Shishir Khetan, Nathan Segall, Joel Solis, Adam Brosz, Carlos Fierro, Howard Schwartz, Kathleen Neuzil, Larry Corey, Peter Gilbert, Holly Janes, Dean Follmann, Mary Marovich, John Mascola, Laura Polakowski, Julie Ledgerwood, Barney S. Graham, Hamilton Bennett, Rolando Pajon, Conor Knightly, Brett Leav, Weiping Deng, Honghong Zhou, Shu Han, Melanie Ivarsson, Jacqueline Miller, Tal Zaks, and COVE Study Group. Efficacy and Safety of the mRNA-1273 SARS-CoV-2 Vaccine. The New England Journal of Medicine, 384(5):403–416, February 2021.

[4] Jeffrey S. Weber, Matteo S. Carlino, Adnan Khattak, Tarek Meniawy, George Ansstas, Matthew H. Taylor, Kevin B. Kim, Meredith McKean, Georgina V. Long, Ryan J. Sullivan, Mark Faries, Thuy T. Tran, C. Lance Cowey, Andrew Pecora, Montaser Shaheen, Jennifer Segar, Theresa Medina, Victoria Atkinson, Geoffrey T. Gibney, Jason J. Luke, Sajeve Thomas, Elizabeth I. Buchbinder, Jane A. Healy, Mo Huang, Manju Morrissey, Igor Feldman, Vasudha Sehgal, Celine Robert-Tissot, Peijie Hou, Lili Zhu, Michelle Brown, Praveen Aanur, Robert S. Meehan, and Tal Zaks. Individualised neoantigen therapy mRNA-4157 (V940) plus pembrolizumab versus pembrolizumab monotherapy in resected melanoma (KEYNOTE-942): a randomised, phase 2b study. The Lancet, 403(10427):632–644, February 2024.

[5] U. Sahin, M. Schmidt, E. Derhovanessian, A. Cortini, I. Vogler, T. Omokoko, E. Godehardt, S. Attig, S. Newrzela, J. Grützner, N. Bidmon, S. Bolte, S. Brachtendorf, T. Stuhlmann, D. Langer, D. Brüne, J. Blake, A. Feldner, H. Lindman, A. Schneeweiss, M. Eichbaum, and Ö Türeci. Individualized mRNA vaccines evoke durable T cell immunity in adjuvant TNBC. Nature, 651(8107):1088–1096, March 2026.

[6] Mirco J. Friedrich, Julie Pham, Jiakun Tian, Hongyu Chen, Jiahao Huang, Niklas Kehl, Sophia Liu, Blake Lash, Fei Chen, Xiao Wang, Rhiannon K. Macrae, and Feng Zhang. Transient hepatic reconstitution of trophic factors enhances aged immunity. Nature, pages 1–9, December 2025.

[7] Gavin Hanson and Jeff Coller. Codon optimality, bias and usage in translation and mRNA decay. Nature reviews. Molecular cell biology, 19(1):20–30, January 2018.

[8] Kimberly A Dittmar, Jeffrey M Goodenbour, and Tao Pan. Tissue-Specific Differences in Human Transfer RNA Expression. PLOS Genetics, 2(12):e221, December 2006.

[9] Hila Gingold, Disa Tehler, Nanna R. Christoffersen, Morten M. Nielsen, Fazila Asmar, Susanne M. Kooistra, Nicolaj S. Christophersen, Lise Lotte Christensen, Michael Borre, Karina D. Sørensen, Lars D. Andersen, Claus L. Andersen, Esther Hulleman, Tom Wurdinger, Elisabeth Ralfkiær, Kristian Helin, Kirsten Grønbæk, Torben Ørntoft, Sebastian M. Waszak, Orna Dahan, Jakob Skou Pedersen, Anders H. Lund, and Yitzhak Pilpel. A Dual Program for Translation Regulation in Cellular Proliferation and Differentiation. Cell, 158(6):1281–1292, September 2014.

[10] Xavier Hernandez-Alias, Hannah Benisty, Leandro G. Radusky, Luis Serrano, and Martin H. Schaefer. Using protein-per-mRNA differences among human tissues in codon optimization. Genome Biology, 24(1):34, February 2023.

[11] Mridu Kapur, Michael J. Molumby, Carlos Guzman, Sven Heinz, and Susan L. Ackerman. Cell-type-specific expression of tRNAs in the brain regulates cellular homeostasis. Neuron, 112(9):1397–1415.e6, May 2024.

[12] P. M. Sharp and W. H. Li. The codon Adaptation Index–a measure of directional synonymous codon usage bias, and its potential applications. Nucleic Acids Research, 15(3):1281–1295, February 1987.

[13] Claes Gustafsson, Sridhar Govindarajan, and Jeremy Minshull. Codon bias and heterologous protein expression. Trends in Biotechnology, 22(7):346–353, July 2004.

[14] Joshua B. Plotkin and Grzegorz Kudla. Synonymous but not the same: the causes and consequences of codon bias. Nature reviews. Genetics, 12(1):32–42, January 2011.

[15] Douglas Meyer, Jacob Kames, Haim Bar, Anton A. Komar, Aikaterini Alexaki, Juan Ibla, Ryan C. Hunt, Luis V. Santana-Quintero, Anton Golikov, Michael DiCuccio, and Chava Kimchi-Sarfaty. Distinct signatures of codon and codon pair usage in 32 primary tumor types in the novel database CancerCoCoPUTs for cancer-specific codon usage. Genome Medicine, 13(1):122, July 2021.

[16] He Zhang, Liang Zhang, Ang Lin, Congcong Xu, Ziyu Li, Kaibo Liu, Boxiang Liu, Xiaopin Ma, Fanfan Zhao, Huiling Jiang, Chunxiu Chen, Haifa Shen, Hangwen Li, David H. Mathews, Yujian Zhang, and Liang Huang. Algorithm for optimized mRNA design improves stability and immunogenicity. Nature, 621(7978):396–403, September 2023.

[17] Zeming Lin, Halil Akin, Roshan Rao, Brian Hie, Zhongkai Zhu, Wenting Lu, Nikita Smetanin, Robert Verkuil, Ori Kabeli, Yaniv Shmueli, Allan dos Santos Costa, Maryam Fazel-Zarandi, Tom Sercu, Salvatore Candido, and Alexander Rives. Evolutionary-scale prediction of atomic level protein structure with a language model, December 2022. Pages: 2022.07.20.500902 Section: New Results.

[18] Thomas Hayes, Roshan Rao, Halil Akin, Nicholas J. Sofroniew, Deniz Oktay, Zeming Lin, Robert Verkuil, Vincent Q. Tran, Jonathan Deaton, Marius Wiggert, Rohil Badkundri, Irhum Shafkat, Jun Gong, Alexander Derry, Raul S. Molina, Neil Thomas, Yousuf A. Khan, Chetan Mishra, Carolyn Kim, Liam J. Bartie, Matthew Nemeth, Patrick D. Hsu, Tom Sercu, Salvatore Candido, and Alexander Rives. Simulating 500 million years of evolution with a language model, December 2024. Pages: 2024.07.01.600583 Section: New Results.

[19] Yikun Zhang, Mei Lang, Jiuhong Jiang, Zhiqiang Gao, Fan Xu, Thomas Litfin, Ke Chen, Jaswinder Singh, Xiansong Huang, Guoli Song, Yonghong Tian, Jian Zhan, Jie Chen, and Yaoqi Zhou. Multiple sequence alignment-based RNA language model and its application to structural inference. Nucleic Acids Research, 52(1):e3–e3, January 2024.

[20] Hugo Dalla-Torre, Liam Gonzalez, Javier Mendoza-Revilla, Nicolas Lopez Carranza, Adam Henryk Grzywaczewski, Francesco Oteri, Christian Dallago, Evan Trop, Bernardo P. de Almeida, Hassan Sirelkhatim, Guillaume Richard, Marcin Skwark, Karim Beguir, Marie Lopez, and Thomas Pierrot. Nucleotide Transformer: building and evaluating robust foundation models for human genomics. Nature Methods, pages 1–11, November 2024.

[21] Eric Nguyen, Michael Poli, Matthew G. Durrant, Brian Kang, Dhruva Katrekar, David B. Li, Liam J. Bartie, Armin W. Thomas, Samuel H. King, Garyk Brixi, Jeremy Sullivan, Madelena Y. Ng, Ashley Lewis, Aaron Lou, Stefano Ermon, Stephen A. Baccus, Tina Hernandez-Boussard, Christopher Ré, Patrick D. Hsu, and Brian L. Hie. Sequence modeling and design from molecular to genome scale with Evo. Science, 386(6723):eado9336, November 2024.

[22] Žiga Avsec, Natasha Latysheva, Jun Cheng, Guido Novati, Kyle R. Taylor, Tom Ward, Clare Bycroft, Lauren Nicolaisen, Eirini Arvaniti, Joshua Pan, Raina Thomas, Vincent Dutordoir, Matteo Perino, Soham De, Alexander Karollus, Adam Gayoso, Toby Sargeant, Anne Mottram, Lai Hong Wong, Pavol Drotár, Adam Kosiorek, Andrew Senior, Richard Tanburn, Taylor Applebaum, Souradeep Basu, Demis Hassabis, and Pushmeet Kohli. Advancing regulatory variant effect prediction with AlphaGenome. Nature, 649(8099):1206–1218, January 2026.

[23] Yanyi Chu, Dan Yu, Yupeng Li, Kaixuan Huang, Yue Shen, L. Cong, Jason Zhang, and Mengdi Wang. A 5 UTR language model for decoding untranslated regions of mRNA and function predictions. Nature Machine Intelligence, 6(4):449–460, April 2024.

[24] Sizhen Li, Saeed Moayedpour, Ruijiang Li, Michael Bailey, Saleh Riahi, Lorenzo Kogler-Anele, Milad Miladi, Jacob Miner, Fabien Pertuy, Dinghai Zheng, Jun Wang, Akshay Balsubramani, Khang Tran, Minnie Zacharia, Monica Wu, Xiaobo Gu, Ryan Clinton, Carla Asquith, Joseph Skaleski, Lianne Boeglin, Sudha Chivukula, Anusha Dias, Tod Strugnell, Fernando Ulloa Montoya, Vikram Agarwal, Ziv Bar-Joseph, and Sven Jager. CodonBERT large language model for mRNA vaccines. Genome Research, 34(7):1027–1035, July 2024.

[25] Binita Rajbanshi and Anuj Guruacharya. codonGPT: reinforcement learning on a generative language model enables scalable mRNA design. Nucleic Acids Research, 53(22):gkaf1345, November 2025.

[26] He Zhang, Hailong Liu, Yushan Xu, Haoran Huang, Yiming Liu, Jia Wang, Yan Qin, Haiyan Wang, Lili Ma, Zhiyuan Xun, Xuzhuang Hou, Timothy K. Lu, and Jicong Cao. Deep generative models design mRNA sequences with enhanced translational capacity and stability. Science, 390(6773):eadr8470, November 2025.

[27] Ying Xiong, Aowen Wang, Yu Kang, Chao Shen, Chang-Yu Hsieh, and Tingjun Hou. mRNABERT: advancing mRNA sequence design with a universal language model and comprehensive dataset. Nature Communications, 16(1):10371, November 2025.

[28] Adibvafa Fallahpour, Vincent Gureghian, Guillaume J. Filion, Ariel B. Lindner, and Amir Pandi. Codon-Transformer: a multispecies codon optimizer using context-aware neural networks, September 2024. Pages: 2024.09.13.612903 Section: New Results.

[29] Long-Kai Huang, Rongyi Zhu, Bing He, and Jianhua Yao. Steering Protein Language Models, September 2025. arXiv:2509.07983 [q-bio].

[30] Junhao Xiong, Ishan Gaur, Maria Lukarska, Hunter Nisonoff, Luke M. Oltrogge, David F. Savage, and Jennifer Listgarten. Property guidance for protein sequence generative models with ProteinGuide. Nature Biotechnology, July 2026.

[31] Albert Gu and Tri Dao. Mamba: Linear-Time Sequence Modeling with Selective State Spaces, May 2024. arXiv:2312.00752.

[32] Carlos Outeiral and Charlotte M. Deane. Codon language embeddings provide strong signals for use in protein engineering. Nature Machine Intelligence, 6(2):170–179, February 2024.

[33] Tao Shen, Zhihang Hu, Siqi Sun, D. Liu, Felix Wong, Jiuming Wang, Jiayang Chen, Yixuan Wang, Liang Hong, Jin Xiao, Liangzhen Zheng, Tejas Krishnamoorthi, Irwin King, Sheng Wang, Peng Yin, James J. Collins, and Yu Li. Accurate RNA 3D structure prediction using a language model-based deep learning approach. Nature Methods, 21(12):2287–2298, December 2024.

[34] Sharrol Bachas, Goran Rakocevic, David Spencer, Anand V. Sastry, Robel Haile, John M. Sutton, George Kasun, Andrew Stachyra, Jahir M. Gutierrez, Edriss Yassine, Borka Medjo, Vincent Blay, Christa Kohnert, Jennifer T. Stanton, Alexander Brown, Nebojsa Tijanic, Cailen McCloskey, Rebecca Viazzo, Rebecca Consbruck, Hayley Carter, Simon Levine, Shaheed Abdulhaqq, Jacob Shaul, Abigail B. Ventura, Randal S. Olson, Engin Yapici, Joshua Meier, Sean McClain, Matthew Weinstock, Gregory Hannum, Ariel Schwartz, Miles Gander, and Roberto Spreafico. Antibody optimization enabled by artificial intelligence predictions of binding affinity and naturalness, August 2022. Pages: 2022.08.16.504181 Section: New Results.

[35] David A. Constant, Jahir M. Gutierrez, Anand V. Sastry, Rebecca Viazzo, Nicholas R. Smith, Jubair Hossain, David A. Spencer, Hayley Carter, Abigail B. Ventura, Michael T. M. Louie, Christa Kohnert, Rebecca Consbruck, Joshua Bennett, Kenneth A. Crawford, John M. Sutton, Anneliese Morrison, Andrea K. Steiger, Kerianne A. Jackson, Jennifer T. Stanton, Shaheed Abdulhaqq, Gregory Hannum, Joshua Meier, Matthew Weinstock, and Miles Gander. Deep learning-based codon optimization with large-scale synonymous variant datasets enables generalized tunable protein expression, February 2023. Pages: 2023.02.11.528149 Section: New Results.

[36] Adibvafa Fallahpour, Vincent Gureghian, Guillaume J. Filion, Ariel B. Lindner, and Amir Pandi. CodonTransformer: a multispecies codon optimizer using context-aware neural networks. Nature Communications, 16(1):3205, April 2025.

[37] Carla Cummins, Alisha Ahamed, Raheela Aslam, Josephine Burgin, Rajkumar Devraj, Ossama Edbali, Dipayan Gupta, Peter W Harrison, Muhammad Haseeb, Sam Holt, Talal Ibrahim, Eugene Ivanov, Suran Jayathilaka, Vishnukumar Kadhirvelu, Simon Kay, Manish Kumar, Ankur Lathi, Rasko Leinonen, Fabio Madeira, Nandana Madhusoodanan, Milena Mansurova, Colman O’Cathail, Matt Pearce, Stéphane Pesant, Nadim Rahman, Jeena Rajan, Gabriele Rinck, Sandeep Selvakumar, Alexey Sokolov, Swati Suman, Ross Thorne, Prabhat Totoo, Senthilnathan Vijayaraja, Zahra Waheed, Ahmad Zyoud, Rodrigo Lopez, Tony Burdett, and Guy Cochrane. The European Nucleotide Archive in 2021. Nucleic Acids Research, 50(D1):D106–D110, January 2022.

[38] Weizhong Li and Adam Godzik. Cd-hit: a fast program for clustering and comparing large sets of protein or nucleotide sequences. Bioinformatics (Oxford, England), 22(13):1658–1659, July 2006. Number: 13.

[39] James Heuschkel, Laura Kingsley, Noah Pefaur, Andrew Nixon, and Steven Cramer. Advancing Codon Language Modeling with Synonymous Codon Constrained Masking, August 2025. ISSN: 2692-8205 Pages: 2025.08.19.671089 Section: New Results.

[40] Alicia A. Bicknell, David W. Reid, Marissa C. Licata, Adriana K. Jones, Yi Min Cheng, Mengying Li, Chiaowen Joyce Hsiao, Christopher S. Pepin, Mihir Metkar, Yevgen Levdansky, Brian R. Fritz, Elizaveta A. Andrianova, Ruchi Jain, Eugene Valkov, Caroline Köhrer, and Melissa J. Moore. Attenuating ribosome load improves protein output from mRNA by limiting translation-dependent mRNA decay. Cell Reports, 43(4), April 2024.

[41] Kathrin Leppek, Gun Woo Byeon, Wipapat Kladwang, Hannah K. Wayment-Steele, Craig H. Kerr, Adele F. Xu, Do Soon Kim, Ved V. Topkar, Christian Choe, Daphna Rothschild, Gerald C. Tiu, Roger Wellington-Oguri, Kotaro Fujii, Eesha Sharma, Andrew M. Watkins, John J. Nicol, Jonathan Romano, Bojan Tunguz, Fernando Diaz, Hui Cai, Pengbo Guo, Jiewei Wu, Fanyu Meng, Shuai Shi, Eterna Participants, Philip R. Dormitzer, Alicia Solórzano, Maria Barna, and Rhiju Das. Combinatorial optimization of mRNA structure, stability, and translation for RNA-based therapeutics. Nature Communications, 13(1):1536, March 2022.

[42] Yair Schiff, Chia Hsiang Kao, Aaron Gokaslan, Tri Dao, Albert Gu, and Volodymyr Kuleshov. Caduceus: Bi-Directional Equivariant Long-Range DNA Sequence Modeling. In Proceedings of the 41st International Conference on Machine Learning, pages 43632–43648. PMLR, July 2024.

[43] Ilya Loshchilov and Frank Hutter. Decoupled Weight Decay Regularization, January 2019. arXiv:1711.05101 [cs].

[44] Laurens van der Maaten and Geoffrey Hinton. Visualizing Data using t-SNE. Journal of Machine Learning Research, 9(86):2579–2605, 2008.

[45] Kaiming He, Xiangyu Zhang, Shaoqing Ren, and Jian Sun. Deep Residual Learning for Image Recognition. In 2016 IEEE Conference on Computer Vision and Pattern Recognition (CVPR), pages 770–778, June 2016. ISSN: 1063-6919.

[46] Alex Wang and Kyunghyun Cho. BERT has a Mouth, and It Must Speak: BERT as a Markov Random Field Language Model, April 2019. arXiv:1902.04094 [cs].

[47] Y. Nakamura, T. Gojobori, and T. Ikemura. Codon usage tabulated from international DNA sequence databases: status for the year 2000. Nucleic Acids Research, 28(1):292, January 2000.

[48] Julian Salazar, Davis Liang, Toan Q. Nguyen, and Katrin Kirchhoff. Masked Language Model Scoring. In Proceedings of the 58th Annual Meeting of the Association for Computational Linguistics, pages 2699–2712, 2020. arXiv:1910.14659 [cs].

