## Supplementary Information for "CodonMamba: a foundation model for programmable mRNA coding sequence design"

#### Supplementary Notes

##### Downstream benchmark datasets and evaluation

CodonMamba was evaluated on 12 public mRNA-related prediction benchmarks spanning protein expression, mRNA stability and degradation, and regulatory phenotypes. Processed datasets were drawn primarily from benchmark collections released with CodonBERT [1], CaLM [2] and SynCodonLM [3]. Protein-expression tasks included mRFP expression and categorical protein expression in *Escherichia coli* [4, 5], fungal protein expression [6], synonymous GFP expression in *E. coli* [7], and expression of influenza haemagglutinin mRNAs in HeLa cells [1]. Stability-related tasks comprised transcript stability across four vertebrate species [8], degradation of SARS-CoV-2 vaccine-related mRNAs [9], and mRNA abundance of synonymous GFP and *TDH3* variants in *Saccharomyces cerevisiae* [10]. Regulatory benchmarks included Tc-riboswitch activity [11], multi-label mRNA subcellular localization [12, 13], and toxicity associated with synonymous GFP variants in *E. coli* [14].

Regression benchmarks were evaluated using Spearman's rank correlation coefficient ($\rho$), multi-class classification using accuracy, and the multi-label subcellular localization benchmark using support-weighted F1 after sigmoid transformation and a probability threshold of 0.5. Dataset sizes, sequence lengths, prediction endpoints, task types and biological sources are summarized in Supplementary Table S1.

##### CDS generation and decoding strategies

Unless otherwise stated, all primary generation analyses in Figs. 4–6 used greedy decoding with a 12-step linear Mask-and-Predict schedule [15], selecting the admissible codon with the highest final logit at each scheduled update. Alternative decoding strategies were evaluated as robustness analyses in Supplementary Fig. S9 while retaining the same synonym-aware constraints and, where applicable, the same host codon usage prior. These included unbiased sampling from the constrained codon distribution ($T = 0.8$), biased sampling with $q(c) \propto p(c)^{\alpha}$ and $\alpha = e^2$ or $e^3$, and beam search with a beam size of 8 and up to three candidate codons per updated position. Beam candidates were ranked by cumulative log-probability within each Mask-and-Predict iteration.

##### CDS design baselines and generation

Baseline CDSs were either obtained directly from the benchmark released with GEMORNA [16] or generated using the publicly available implementations and pretrained models of the respective methods. All comparisons used the same target-protein benchmark as the CodonMamba generation analyses.

Natural CDSs, random synonymous controls, GEMORNA-generated CDSs, human-reference CAI-optimized CDSs and human LinearDesign designs were obtained directly from the GEMORNA benchmark [16] and analysed without regeneration. For cross-host analyses, corresponding CAI-optimized and LinearDesign [17] sequences for Mouse, *E. coli* and Yeast were generated separately using the codon usage reference for each species.

codonGPT [18] and CodonTransformer [19] CDSs were generated for the same target proteins using the publicly released source code and pretrained models provided by the respective authors. Generation followed the conditioning options supported by each original implementation.

For the cross-host comparison, CodonTransformer was generated using the corresponding supported species condition. codonGPT did not use an interchangeable host-specific inference-time condition in the configuration evaluated here; therefore, a single codonGPT-generated CDS for each target protein was evaluated against each host codon usage reference. GEMORNA sequences were likewise treated as fixed benchmark outputs and evaluated against each host reference without regeneration.

##### Evaluation of generated CDSs

Unless otherwise stated, CDSs were converted to the RNA alphabet, tokenized into non-overlapping codons and stripped of incomplete terminal triplets before analysis. Stop codons were excluded from codon-usage and host-matching metrics. For paired comparisons, sequences were restricted to their shared codon-aligned length where required.

**Sequence divergence and codon-usage similarity.** Divergence from the natural CDS was quantified using normalized codon Hamming distance (NHD). For two aligned codon sequences $c = (c_1, \ldots, c_L)$ and $c' = (c'_1, \ldots, c'_L)$,

$$\mathrm{NHD}(c, c') = \frac{1}{L} \sum_{i=1}^{L} \mathbb{I}[c_i \neq c'_i]. \tag{1}$$

Codon match rate was defined as $1 - \mathrm{NHD}$. Synonymous substitution number was the number of differing codon positions that retained the same non-stop amino acid.

Gene-level codon-usage similarity was calculated as the cosine similarity between 61-dimensional sense-codon count vectors:

$$\mathrm{CUS}(c, c') = \frac{\mathbf{v}(c) \cdot \mathbf{v}(c')}{\|\mathbf{v}(c)\| \, \|\mathbf{v}(c')\| + \varepsilon}, \tag{2}$$

where $\mathbf{v}$ contains counts of the 61 sense codons in a fixed order and $\varepsilon = 10^{-12}$. The same 61-dimensional representation was used for cross-host codon-usage profile comparisons.

**Host codon matching.** Species-level codon usage frequencies for Human, Mouse, *E. coli* and Yeast were obtained from the Codon Usage Database (Kazusa) [20]. For codon $c$ encoding amino acid $a$, relative adaptiveness was defined within each synonymous codon family as

$$w(c) = \frac{f(c)}{\max_{c' \in \mathscr{V}(a)} f(c')}, \tag{3}$$

where $f(c)$ is the corresponding species-level codon usage frequency.

Codon adaptation index (CAI) was calculated as the geometric mean of these relative adaptiveness weights across scored sense codons:

$$\mathrm{CAI}(S) = \exp\left[\frac{1}{L_s} \sum_{k=1}^{L_s} \log w(c_k)\right], \tag{4}$$

where $L_s$ denotes the number of scored codons [21, 22, 23].

To provide a complementary transcript-level measure of synonymous codon preference, we calculated scaled relative synonymous codon usage (sRSCU). Scaling conventional RSCU by the maximum value within each synonymous family is equivalent to the relative adaptiveness weight above, $\mathrm{sRSCU}(c) = w(c)$. Transcript-level sRSCU was therefore defined as

$$\mathrm{sRSCU}_{\mathrm{gene}}(S) = \frac{1}{L_s} \sum_{k=1}^{L_s} w(c_k). \tag{5}$$

Thus, CAI summarizes host matching through a geometric mean, whereas sRSCU uses the corresponding arithmetic mean.

**Nucleotide composition and RNA structure.** GC content and uridine percentage were calculated as the fractions of nucleotides corresponding to G or C and to U, respectively. RNA minimum free energy (MFE) was predicted using RNAfold from the ViennaRNA package [24] and normalized by nucleotide length:

$$\mathrm{nMFE}(S) = \frac{\mathrm{MFE}(S)}{n}, \tag{6}$$

where $n$ is the CDS length in nucleotides.

**Local codon-context features.** Rare codons were defined using the host-specific relative adaptiveness weights above, with $w(c) < 0.20$ classified as rare. Rare-codon density was calculated as the number of rare codons divided by CDS nucleotide length.

Unwanted codon pairs were defined using a CoCoPUTs-style human codon-pair usage reference (*Homo sapiens*, taxid 9606) [25]. Codon pairs within the lowest 5% of the reference score distribution were classified as unwanted, and their density was normalized by CDS nucleotide length.

Potential frameshift-associated slippery sites were identified from predefined candidate $-1$ slippery heptamers and $+1$ frameshift-related sequence contexts, including homopolymeric runs and rare-codon-associated motifs [26, 27]. Candidate positions were evaluated in the annotated coding frame, with rare-codon triggering defined using the same $w(c) < 0.20$ threshold. Slippery-site density was calculated as the number of unique candidate positions divided by CDS nucleotide length.

#### Additional analyses of inference-time host priors

**Host-free decoding ablation ($\beta = 0$).** To isolate the contribution of the inference-time host prior, we generated a single host-free CDS set at $\beta = 0$ and evaluated the same sequences against the Human, Mouse, *E. coli* and Yeast codon usage references. Median CAI values were 0.828, 0.822, 0.672 and 0.463, respectively (Supplementary Table S4). Replacing only the codon usage prior at $\beta = 2.0$ increased the corresponding matched-host CAI, with particularly large shifts for *E. coli* (0.672 to 0.881) and Yeast (0.463 to 0.980; Fig. 6B). Thus, the diagonal host-matching pattern observed under prior-guided decoding is attributable to inference-time prior switching, rather than host-specific modification of the pretrained CodonMamba backbone.

#### Supplementary Figures

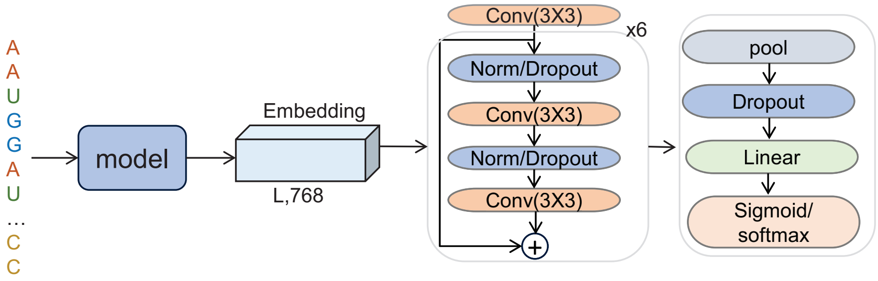


Supplementary Fig. S1: **Downstream evaluation with a lightweight prediction head.** Pretrained sequence representations of size $L \times d$ are processed by a shared ResNet-style convolutional head comprising residual blocks [28], normalization, dropout and pooling, followed by a task-specific linear output layer for regression, multi-class classification or multi-label classification.

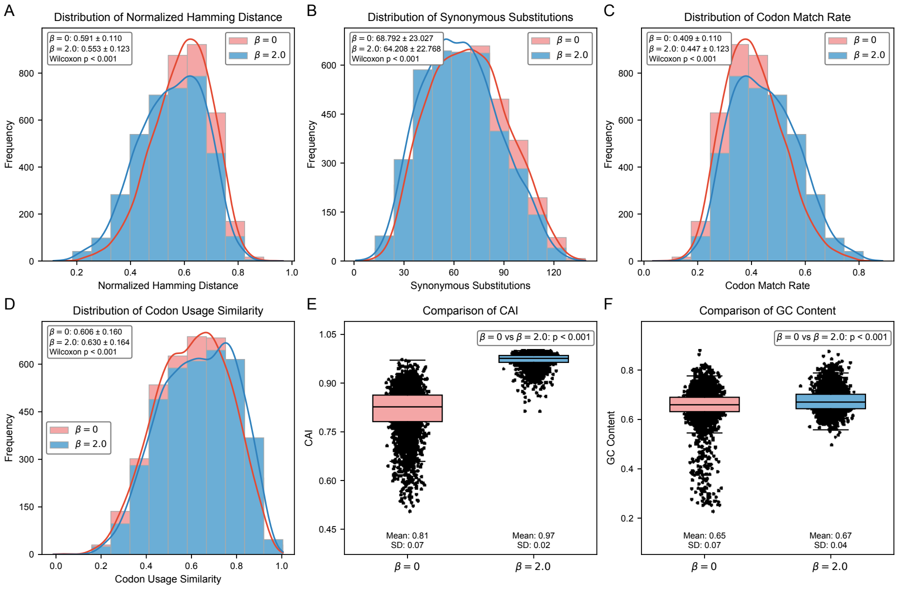


Supplementary Fig. S2: **Effect of codon usage prior on synonymous CDS generation.** CodonMamba sequences generated without a host codon usage prior ($\beta = 0$) were compared with sequences generated using the human prior ($\beta = 2.0$); synonym-aware logit masking was applied in both conditions. **A–D**, Natural-referenced normalized codon Hamming distance, synonymous substitution count, codon match rate and codon-usage similarity. **E**, Human codon adaptation index (CAI). **F**, GC content. Paired comparisons were assessed using two-sided Wilcoxon signed-rank tests.

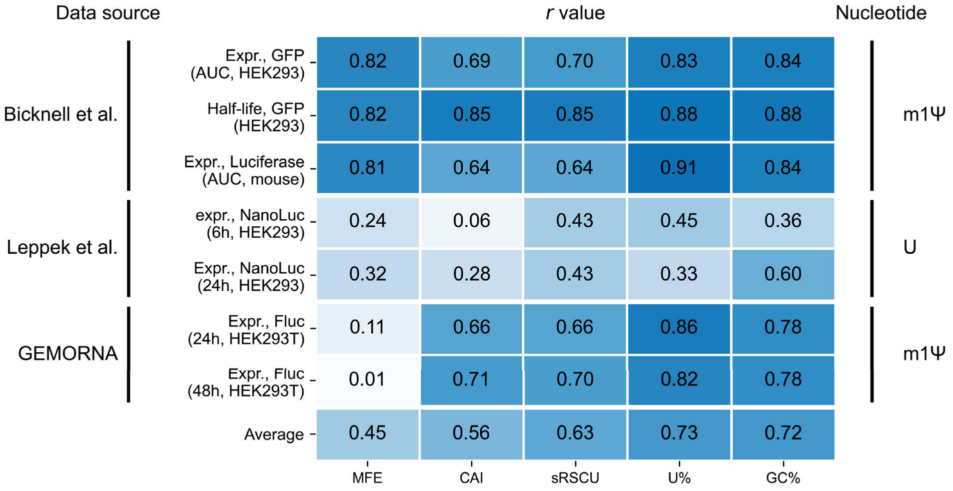


Supplementary Fig. S3: **Associations between CDS features and experimentally measured mRNA performance.** Heatmap shows absolute Pearson correlation coefficients ($|r|$) between five sequence features and experimental protein expression or mRNA stability measurements across three public datasets. Rows denote individual experimental conditions, and the final row shows the mean $|r|$ across readouts. Feature values were obtained from the public benchmark data [16], except for sRSCU, which was calculated in this study.

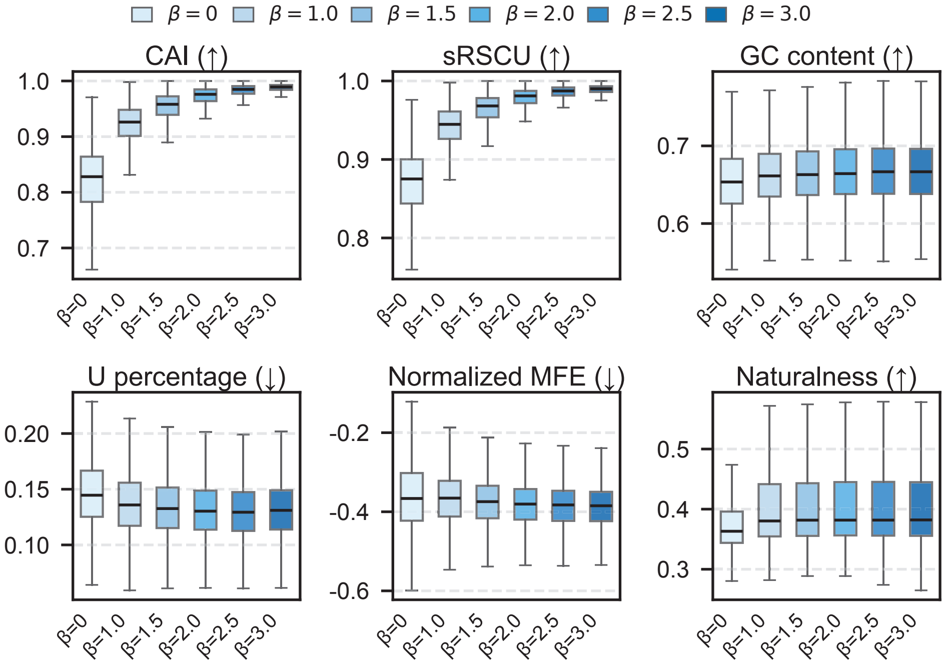


Supplementary Fig. S4: **Effect of host-prior strength on CodonMamba-generated sequence properties.** Boxplots show six sequence properties for CDSs generated with $\beta = 0$–3.0, including CAI, sRSCU, GC content, uridine percentage, normalized minimum free energy and CodonMamba-derived naturalness.

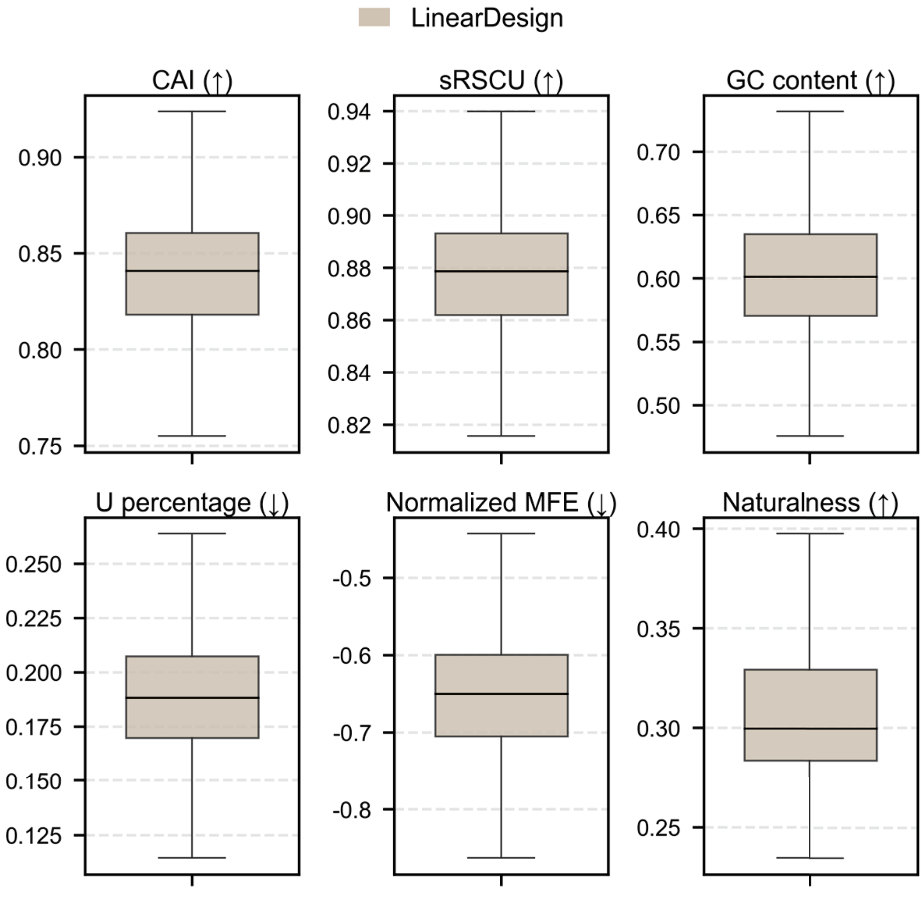


Supplementary Fig. S5: **Sequence-property profiles of LinearDesign CDSs.** Boxplots summarize six sequence properties for LinearDesign-generated CDSs obtained from the GEMORNA benchmark.

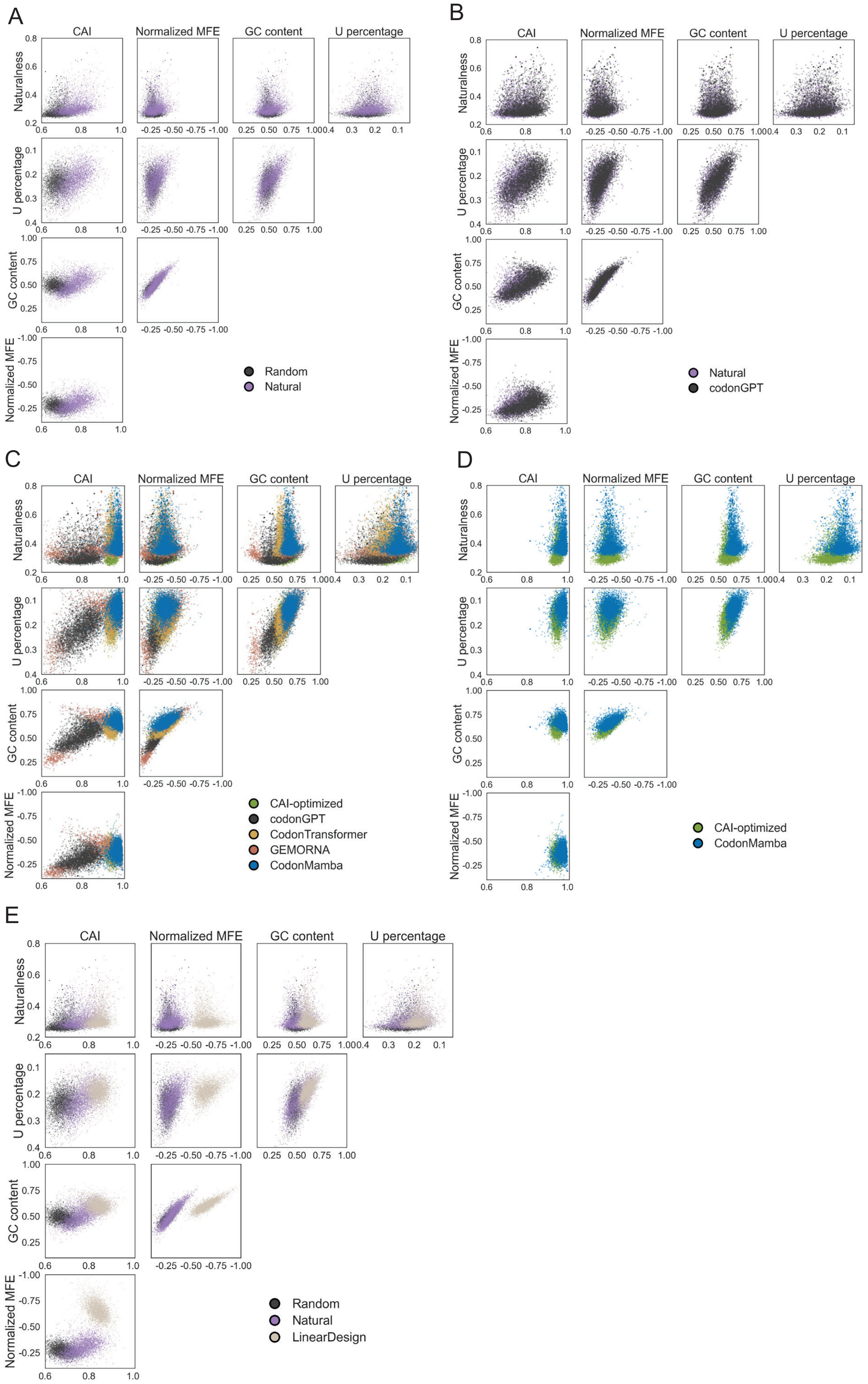


Supplementary Fig. S6: **Pairwise distributions of sequence properties across CDS design methods. A–E**, Pairwise comparisons of selected CDS properties across natural, random and computationally designed sequence groups.

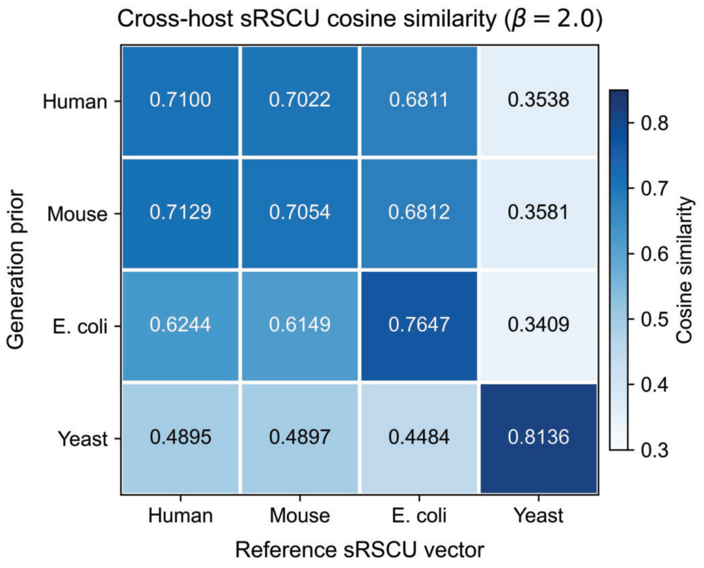


Supplementary Fig. S7: **Cross-host similarity of codon-usage profiles.** Cosine similarity between the mean 61-dimensional sense-codon usage profile of CDSs generated with each host prior (rows) and the corresponding species-specific codon usage references (columns). Diagonal enrichment indicates preferential matching to the specified host.

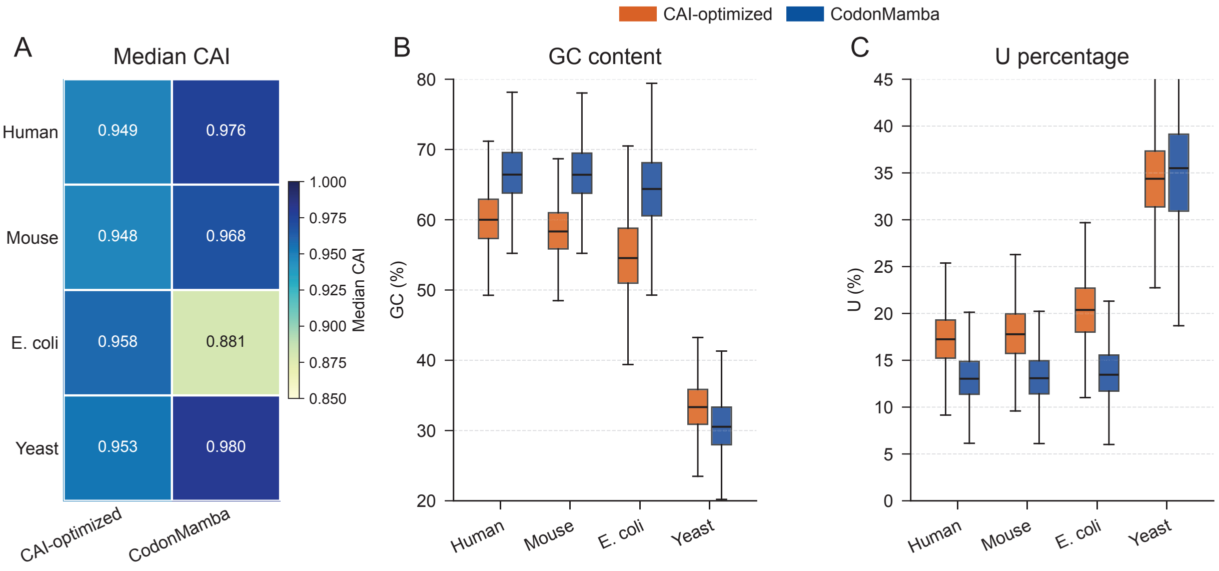


Supplementary Fig. S8: **Comparison of CodonMamba with CAI-optimized designs.** **A**, Median CAI of CAI-optimized and CodonMamba-generated CDSs under matched host references. **B,C**, GC content and uridine percentage. The human CAI-optimized sequences were obtained from the GEMORNA benchmark, whereas the Mouse, *E. coli* and Yeast controls were generated using the corresponding species-specific codon usage references. CAI-optimized sequences are shown as a positive control because high CAI was explicitly favoured during their construction.

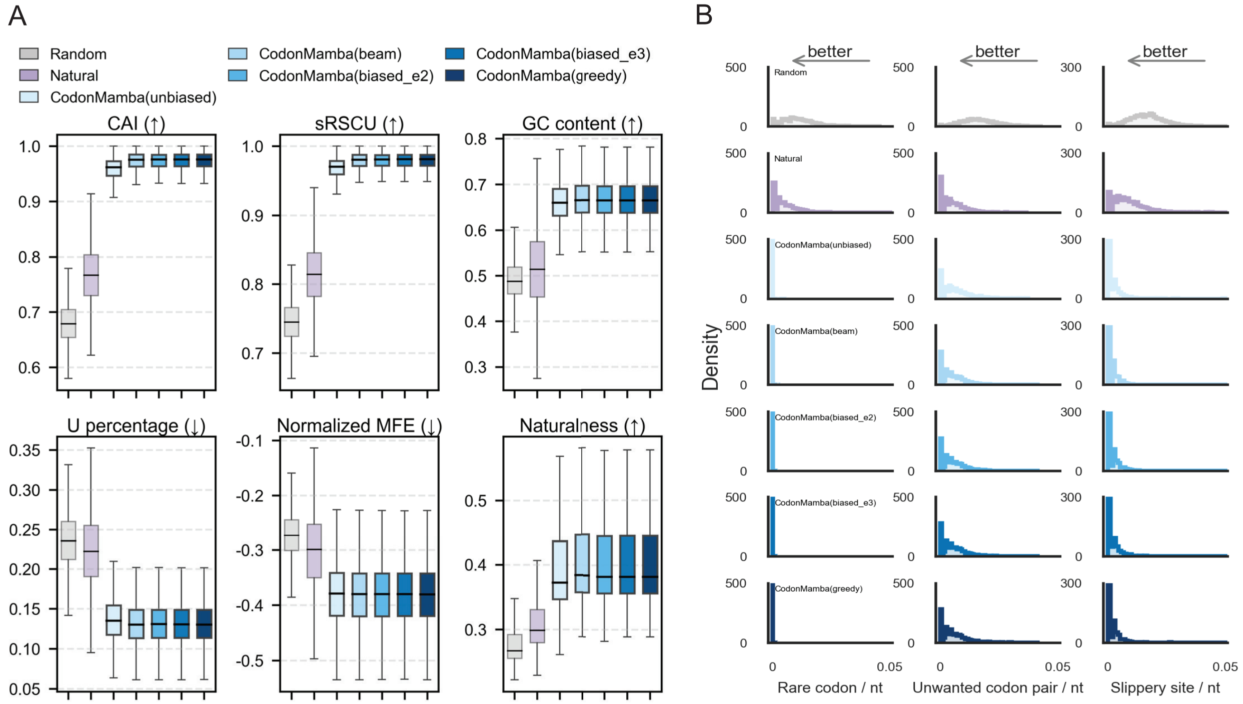


Supplementary Fig. S9: **Robustness of generated CDS properties to decoding strategy. A**, Six sequence properties of CodonMamba-generated CDSs obtained using unbiased sampling, biased sampling with $\alpha = e^2$ or $e^3$, beam search and greedy decoding, with natural CDSs and random synonymous sequences shown as references. **B**, Rare-codon, unwanted codon-pair and slippery-site densities for the corresponding generated CDSs and reference sequences.

### Supplementary Tables

Supplementary Table S1: Overview of datasets used for downstream evaluation, including prediction target, task type, dataset size, sequence length and biological source.

| Dataset | Target | Task | No. of mRNAs | Sequence length | Species/source |
|---|---|---|---|---|---|
| MLOS Flu Vaccines [1] | Expression | Regression | 167 | 1701–1704 | Influenza |
| Tc-Riboswitches [11] | Switching factor | Regression | 355 | 66–75 | Yeast |
| mRNA Stability [8] | Stability | Regression | 65356 | 30–3066 | human/mouse/frog/fish |
| mRFP Expression [4] | Expression | Regression | 1459 | 678 | *Escherichia coli* |
| *E. coli* Proteins [5] | Expression | Classification | 6348 | 171–3000 | *Escherichia coli* |
| Fungal Expression [6] | Expression | Regression | 7089 | 150–3000 | Fungi |
| SARS-CoV-2 [9] | Degradation | Regression | 2400 | 81 | SARS-CoV-2 |
| Subcellular Localization [2] | Localization | Multi-label classification | 5969 | 72–14910 | multispecies |
| mRNA Toxicity [14] | Toxicity | Regression | 97 | 720 | *Escherichia coli* |
| GFP Expression [7] | Expression | Regression | 219 | 30 | *Escherichia coli* |
| GFP mRNA Abundance [10] | Abundance | Regression | 2432 | 36 | Yeast |
| TDH3 mRNA Abundance [10] | Abundance | Regression | 523 | 36 | Yeast |

*Note:* Original author-provided training, validation and test partitions were retained when available. Datasets without predefined partitions were split at a 0.70/0.15/0.15 ratio. All models were evaluated on identical partitions within each benchmark.

Supplementary Table S2: Pretrained baseline models used in downstream benchmarking. One-hot encoding was included separately as a non-pretrained baseline.

| Method | Data type | Context | Token | Architecture | Objective | Parameters | Weights | Code | Pretraining data |
|---|---|---|---|---|---|---|---|---|---|
| mRNABERT [29] | mRNA | 1024 | Codon | Transformer | MLM | 113M | HuggingFace | GitHub | $\sim$18M |
| codonGPT [18] | mRNA | 1024 | Codon | Transformer | NTP | $\sim$0.34M | HuggingFace | GitHub | 3.4M |
| GEMORNA [16] | CDS | – | Codon | Transformer | NTP | 4.4M | GitHub | GitHub | $>$ 1M |
| Helix-mRNA [30] | mRNA | 1024 | Base | Mamba | NTP | 5.19M | HuggingFace | GitHub | 27M |
| CodonBERT [1] | mRNA | 512 | Codon | Transformer | MLM/STP | 87M | GitHub | GitHub | 10M |
| CaLM [2] | mRNA | 1024 | Codon | Transformer | MLM | 87M | CaLM | GitHub | 10M |
| CodonTransformer [19] | CDS | 2048 | Codon | Transformer | MLM | 90M | HuggingFace | GitHub | 1M |
| SpliceBERT [31] | pre-mRNA | 1024 | Base | Transformer | MLM | 20M | Zenodo | GitHub | 2M |
| ESM2 [32] | Protein | 1024 | AA | Transformer | MLM | 650M | GitHub | GitHub | 250M |
| mRNAFM [33] | mRNA | 1024 | Base | Transformer | MLM | 239M | HuggingFace | GitHub | 40M |
| NT [34] | DNA | 6–12K | 6-mer | Transformer | MLM | 50M–2.5B | HuggingFace | GitHub | 174B |

*Note:* MLM, masked language modelling; NTP, next-token prediction; STP, sequence taxonomy prediction; AA, amino acid.

Supplementary Table S3: Performance comparison across 12 mRNA-related benchmark datasets. Accuracy is reported for the *E. coli* protein-expression dataset, weighted F1 score for subcellular localization, and Spearman's rank correlation for all remaining datasets. Values are mean ± s.d. across three independent runs.

| Dataset | OneHot | ESM2 | mRNAFM | SpliceBERT | NT | CodonBERT | Helix-mRNA | CodonTransformer | CaLM | GEMORNA | codonGPT | mRNABERT | CodonMamba |
|---|---|---|---|---|---|---|---|---|---|---|---|---|---|
| mRFP expression | 0.765±0.0007 | 0.000±0.0000 | 0.781±0.0009 | 0.781±0.0001 | 0.793±0.0004 | 0.789±0.0014 | 0.786±0.0000 | 0.827±0.0000 | 0.831±0.0005 | 0.640±0.0001 | 0.754±0.0001 | 0.800±0.0003 | 0.855±0.0004 |
| Fungal expression | 0.697±0.0005 | 0.708±0.0014 | 0.769±0.0015 | 0.672±0.0006 | 0.684±0.0002 | 0.750±0.0001 | 0.708±0.0001 | 0.773±0.0009 | 0.784±0.0006 | 0.670±0.0001 | 0.742±0.0003 | 0.792±0.0013 | 0.821±0.0000 |
| GFP expression | 0.748±0.0014 | 0.129±0.0008 | 0.745±0.0003 | 0.771±0.0004 | 0.718±0.0046 | 0.760±0.0001 | 0.776±0.0010 | 0.787±0.0022 | 0.763±0.0002 | 0.050±0.0298 | 0.797±0.0007 | 0.800±0.0013 | 0.803±0.0010 |
| *E. coli* proteins | 0.430±0.0010 | 0.496±0.0010 | 0.473±0.0030 | 0.498±0.0000 | 0.410±0.0010 | 0.479±0.0010 | 0.477±0.0000 | 0.498±0.0040 | 0.510±0.0000 | 0.490±0.0010 | 0.543±0.0001 | 0.457±0.0027 | 0.531±0.0000 |
| MLOS flu vaccines | 0.247±0.0004 | 0.369±0.0032 | 0.427±0.0180 | 0.239±0.0009 | 0.433±0.0045 | 0.334±0.0117 | 0.454±0.0070 | 0.386±0.0026 | 0.429±0.0099 | 0.307±0.0045 | 0.302±0.0223 | 0.405±0.0009 | 0.510±0.0024 |
| Tc-riboswitches | 0.579±0.0052 | 0.546±0.0017 | 0.521±0.0034 | 0.519±0.0011 | 0.575±0.0078 | 0.572±0.0023 | 0.563±0.0026 | 0.505±0.0062 | 0.577±0.0003 | 0.521±0.0009 | 0.496±0.0013 | 0.521±0.0006 | 0.598±0.0003 |
| SARS-CoV-2 vaccine degradation | 0.769±0.0003 | 0.773±0.0004 | 0.742±0.0001 | 0.806±0.0002 | 0.786±0.0000 | 0.799±0.0004 | 0.770±0.0000 | 0.785±0.0013 | 0.786±0.0000 | 0.778±0.0001 | 0.773±0.0010 | 0.758±0.0007 | 0.809±0.0001 |
| mRNA stability | 0.482±0.0000 | 0.514±0.0000 | 0.518±0.0001 | 0.510±0.0000 | 0.505±0.0001 | 0.510±0.0000 | 0.433±0.0000 | 0.520±0.0000 | 0.534±0.0000 | 0.444±0.0001 | 0.492±0.0003 | 0.533±0.0000 | 0.541±0.0000 |
| Subcellular localization | 0.375±0.0050 | 0.562±0.0004 | 0.672±0.0000 | 0.591±0.0002 | 0.581±0.0003 | 0.639±0.0001 | 0.446±0.0013 | 0.544±0.0001 | 0.736±0.0002 | 0.472±0.0001 | 0.627±0.0002 | 0.680±0.0004 | 0.728±0.0000 |
| TDH3 mRNA abundance | 0.728±0.0016 | 0.269±0.0006 | 0.720±0.0003 | 0.658±0.0035 | 0.723±0.0002 | 0.690±0.0017 | 0.731±0.0046 | 0.502±0.0002 | 0.732±0.0008 | 0.713±0.0007 | 0.735±0.0002 | 0.660±0.0014 | 0.797±0.0001 |
| GFP mRNA abundance | 0.842±0.0000 | 0.453±0.0002 | 0.841±0.0000 | 0.827±0.0002 | 0.821±0.0002 | 0.806±0.0002 | 0.840±0.0001 | 0.710±0.0003 | 0.838±0.0001 | 0.827±0.0000 | 0.841±0.0000 | 0.839±0.0000 | 0.843±0.0000 |
| mRNA toxicity | 0.659±0.0159 | 0.421±0.0209 | 0.489±0.0371 | 0.000±0.0000 | 0.529±0.0295 | 0.571±0.0010 | 0.535±0.0069 | 0.556±0.0005 | 0.545±0.0028 | 0.597±0.0021 | 0.353±0.0167 | 0.549±0.0068 | 0.668±0.0025 |

*Note:* Bold values indicate the best-performing method and italic values the second-best method for each benchmark. All reported metrics are higher-is-better. ESM2 receives identical
93 amino-acid sequences for synonymous CDS variants and therefore does not retain codon-level information.

Supplementary Table S4: CAI of CodonMamba sequences generated without a codon usage prior ($\beta = 0$) and evaluated against four host-specific references ($n = 3{,}765$).

| Evaluation reference | Median CAI | [Q1, Q3] |
|---|---|---|
| Human | 0.828 | [0.783, 0.864] |
| Mouse | 0.822 | [0.771, 0.861] |
| *E. coli* | 0.672 | [0.627, 0.712] |
| Yeast | 0.463 | [0.437, 0.490] |

### References


[1] Sizhen Li, Saeed Moayedpour, Ruijiang Li, Michael Bailey, Saleh Riahi, Lorenzo Kogler-Anele, Milad Miladi, Jacob Miner, Fabien Pertuy, Dinghai Zheng, Jun Wang, Akshay Balsubramani, Khang Tran, Minnie Zacharia, Monica Wu, Xiaobo Gu, Ryan Clinton, Carla Asquith, Joseph Skaleski, Lianne Boeglin, Sudha Chivukula, Anusha Dias, Tod Strugnell, Fernando Ulloa Montoya, Vikram Agarwal, Ziv Bar-Joseph, and Sven Jager. CodonBERT large language model for mRNA vaccines. *Genome Research*, 34(7):1027–1035, July 2024.

[2] Carlos Outeiral and Charlotte M. Deane. Codon language embeddings provide strong signals for use in protein engineering. *Nature Machine Intelligence*, 6(2):170–179, February 2024.

[3] James Heuschkel, Laura Kingsley, Noah Pefaur, Andrew Nixon, and Steven Cramer. Advancing Codon Language Modeling with Synonymous Codon Constrained Masking, August 2025. ISSN: 2692-8205 Pages: 2025.08.19.671089 Section: New Results.

[4] Thijs Nieuwkoop, Barbara R. Terlouw, Katherine G. Stevens, Richard A. Scheltema, Dick de Ridder, John van der Oost, and Nico J. Claassens. Revealing determinants of translation efficiency via whole-gene codon randomization and machine learning. *Nucleic Acids Research*, 51(5):2363–2376, March 2023.

[5] Zundan Ding, Feifei Guan, Guoshun Xu, Yuchen Wang, Yaru Yan, Wei Zhang, Ningfeng Wu, Bin Yao, Huoqing Huang, Tamir Tuller, and Jian Tian. MPEPE, a predictive approach to improve protein expression in *E. coli* based on deep learning. *Computational and Structural Biotechnology Journal*, 20:1142–1153, January 2022.

[6] Rhondene Wint, Asaf Salamov, and Igor V Grigoriev. Kingdom-Wide Analysis of Fungal Protein-Coding and tRNA Genes Reveals Conserved Patterns of Adaptive Evolution. *Molecular Biology and Evolution*, 39(2):msab372, February 2022.

[7] Alexander Schmitz and Fuzhong Zhang. Massively parallel gene expression variation measurement of a synonymous codon library. *BMC Genomics*, 22(1):149, March 2021.

[8] Michay Diez, Santiago Gerardo Medina-Muñoz, Luciana Andrea Castellano, Gabriel Da Silva Pescador, Qiushuang Wu, and Ariel Alejandro Bazzini. iCodon customizes gene expression based on the codon composition. *Scientific Reports*, 12(1):12126, July 2022.

[9] Hannah K. Wayment-Steele, Wipapat Kladwang, Andrew M. Watkins, Do Soon Kim, Bojan Tunguz, Walter Reade, Maggie Demkin, Jonathan Romano, Roger Wellington-Oguri, John J. Nicol, Jiayang Gao, Kazuki Onodera, Kazuki Fujikawa, Hanfei Mao, Gilles Vandewiele, Michele Tinti, Bram Steenwinckel, Takuya Ito, Taiga Noumi, Shujun He, Keiichiro Ishi, Youhan Lee, Fatih Öztürk, King Yuen Chiu, Emin Öztürk, Karim Amer, Mohamed Fares, Eterna Participants, and Rhiju Das. Deep learning models for predicting RNA degradation via dual crowdsourcing. *Nature Machine Intelligence*, 4(12):1174–1184, December 2022.

[10] Siyu Chen, Ke Li, Wenqing Cao, Jia Wang, Tong Zhao, Qing Huan, Yu-Fei Yang, Shaohuan Wu, and Wenfeng Qian. Codon-Resolution Analysis Reveals a Direct and Context-Dependent Impact of Individual Synonymous Mutations on mRNA Level. *Molecular Biology and Evolution*, 34(11):2944–2958, November 2017.

[11] Ann-Christin Groher, Sven Jager, Christopher Schneider, Florian Groher, Kay Hamacher, and Beatrix Suess. Tuning the Performance of Synthetic Riboswitches using Machine Learning. *ACS Synthetic Biology*, 8(1):34–44, January 2019.

[12] Vineet Thumuluri, José Juan Almagro Armenteros, Alexander Rosenberg Johansen, Henrik Nielsen, and Ole Winther. DeepLoc 2.0: multi-label subcellular localization prediction using protein language models. *Nucleic Acids Research*, 50(W1):W228–W234, July 2022.

[13] Christian Dallago, Jody Mou, Kadina E. Johnston, Bruce J. Wittmann, Nicholas Bhattacharya, Samuel Goldman, Ali Madani, and Kevin K. Yang. FLIP: Benchmark tasks in fitness landscape inference for proteins, November 2021. Pages: 2021.11.09.467890 Section: New Results.

[14] Pragya Mittal, James Brindle, Julie Stephen, Joshua B. Plotkin, and Grzegorz Kudla. Codon usage influences fitness through RNA toxicity. *Proceedings of the National Academy of Sciences of the United States of America*, 115(34):8639–8644, August 2018.

[15] Marjan Ghazvininejad, Omer Levy, Yinhan Liu, and Luke Zettlemoyer. Mask-Predict: Parallel Decoding of Conditional Masked Language Models. In *Proceedings of the 2019 Conference on Empirical Methods in Natural Language Processing and the 9th International Joint Conference on Natural Language Processing (EMNLP-IJCNLP)*, pages 6111–6120, Hong Kong, China, 2019. Association for Computational Linguistics.

[16] He Zhang, Hailong Liu, Yushan Xu, Haoran Huang, Yiming Liu, Jia Wang, Yan Qin, Haiyan Wang, Lili Ma, Zhiyuan Xun, Xuzhuang Hou, Timothy K. Lu, and Jicong Cao. Deep generative models design mRNA sequences with enhanced translational capacity and stability. *Science*, 390(6773):eadr8470, November 2025.

[17] He Zhang, Liang Zhang, Ang Lin, Congcong Xu, Ziyu Li, Kaibo Liu, Boxiang Liu, Xiaopin Ma, Fanfan Zhao, Huiling Jiang, Chunxiu Chen, Haifa Shen, Hangwen Li, David H. Mathews, Yujian Zhang, and Liang Huang. Algorithm for optimized mRNA design improves stability and immunogenicity. *Nature*, 621(7978):396–403, September 2023.

[18] Binita Rajbanshi and Anuj Guruacharya. codonGPT: reinforcement learning on a generative language model enables scalable mRNA design. *Nucleic Acids Research*, 53(22):gkaf1345, November 2025.

[19] Adibvafa Fallahpour, Vincent Gureghian, Guillaume J. Filion, Ariel B. Lindner, and Amir Pandi. Codon-Transformer: a multispecies codon optimizer using context-aware neural networks, September 2024. Pages: 2024.09.13.612903 Section: New Results.

[20] Y. Nakamura, T. Gojobori, and T. Ikemura. Codon usage tabulated from international DNA sequence databases: status for the year 2000. *Nucleic Acids Research*, 28(1):292, January 2000.

[21] P M Sharp and W H Li. The codon Adaptation Index–a measure of directional synonymous codon usage bias, and its potential applications. *Nucleic Acids Research*, 15(3):1281–1295, February 1987.

[22] Joshua B. Plotkin and Grzegorz Kudla. Synonymous but not the same: the causes and consequences of codon bias. *Nature reviews. Genetics*, 12(1):32–42, January 2011.

[23] Gavin Hanson and Jeff Coller. Codon optimality, bias and usage in translation and mRNA decay. *Nature Reviews. Molecular Cell Biology*, 19(1):20–30, January 2018.

[24] Ronny Lorenz, Stephan H. Bernhart, Christian Höner zu Siederdissen, Hakim Tafer, Christoph Flamm, Peter F. Stadler, and Ivo L. Hofacker. ViennaRNA Package 2.0. *Algorithms for Molecular Biology*, 6(1):26, November 2011.

[25] Aikaterini Alexaki, Jacob Kames, David D. Holcomb, John Athey, Luis V. Santana-Quintero, Phuc Vihn Nguyen Lam, Nobuko Hamasaki-Katagiri, Ekaterina Osipova, Vahan Simonyan, Haim Bar, Anton A. Komar, and Chava Kimchi-Sarfaty. Codon and Codon-Pair Usage Tables (CoCoPUTs): Facilitating Genetic Variation Analyses and Recombinant Gene Design. *Journal of Molecular Biology*, 431(13):2434–2441, June 2019.

[26] Kyusik Q. Kim, Bhagyashri D. Burgute, Shin-Cheng Tzeng, Crystal Jing, Courtney Jungers, Junya Zhang, Liewei L. Yan, Richard D. Vierstra, Sergej Djuranovic, Bradley S. Evans, and Hani S. Zaher. N1-methylpseudouridine found within COVID-19 mRNA vaccines produces faithful protein products. *Cell Reports*, 40(9):111300, August 2022.

[27] Thomas E. Mulroney, Tuija Pöyry, Juan Carlos Yam-Puc, Maria Rust, Robert F. Harvey, Lajos Kalmar, Emily Horner, Lucy Booth, Alexander P. Ferreira, Mark Stoneley, Ritwick Sawarkar, Alexander J. Mentzer, Kathryn S. Lilley, C. Mark Smales, Tobias von der Haar, Lance Turtle, Susanna Dunachie, Paul Klenerman, James E. D. Thaventhiran, and Anne E. Willis. N1-methylpseudouridylation of mRNA causes +1 ribosomal frameshifting. *Nature*, 625(7993):189–194, January 2024.

[28] Kaiming He, Xiangyu Zhang, Shaoqing Ren, and Jian Sun. Deep Residual Learning for Image Recognition. In *2016 IEEE Conference on Computer Vision and Pattern Recognition (CVPR)*, pages 770–778, June 2016. ISSN: 1063-6919.

[29] Ying Xiong, Aowen Wang, Yu Kang, Chao Shen, Chang-Yu Hsieh, and Tingjun Hou. mRNABERT: advancing mRNA sequence design with a universal language model and comprehensive dataset. *Nature Communications*, 16(1):10371, November 2025.

[30] Matthew Wood, Mathieu Klop, and Maxime Allard. Helix-mRNA: A Hybrid Foundation Model For Full Sequence mRNA Therapeutics, March 2025. arXiv:2502.13785 [q-bio].

[31] Ken Chen, Yue Zhou, Maolin Ding, Yu Wang, Zhixiang Ren, and Yuedong Yang. Self-supervised learning on millions of primary RNA sequences from 72 vertebrates improves sequence-based RNA splicing prediction. *Briefings in Bioinformatics*, 25(3):bbae163, April 2024.

[32] Zeming Lin, Halil Akin, Roshan Rao, Brian Hie, Zhongkai Zhu, Wenting Lu, Nikita Smetanin, Robert Verkuil, Ori Kabeli, Yaniv Shmueli, Allan Dos Santos Costa, Maryam Fazel-Zarandi, Tom Sercu, Salvatore Candido, and Alexander Rives. Evolutionary-scale prediction of atomic-level protein structure with a language model. *Science (New York, N.Y.)*, 379(6637):1123–1130, March 2023.

[33] Tao Shen, Zhihang Hu, Siqi Sun, Di Liu, Felix Wong, Jiuming Wang, Jiayang Chen, Yixuan Wang, Liang Hong, Jin Xiao, Liangzhen Zheng, Tejas Krishnamoorthi, Irwin King, Sheng Wang, Peng Yin, James J. Collins, and Yu Li. Accurate RNA 3D structure prediction using a language model-based deep learning approach. *Nature Methods*, 21(12):2287–2298, December 2024.

[34] Hugo Dalla-Torre, Liam Gonzalez, Javier Mendoza-Revilla, Nicolas Lopez Carranza, Adam Henryk Grzywaczewski, Francesco Oteri, Christian Dallago, Evan Trop, Bernardo P. de Almeida, Hassan Sirelkhatim, Guillaume Richard, Marcin Skwark, Karim Beguir, Marie Lopez, and Thomas Pierrot. Nucleotide Transformer: building and evaluating robust foundation models for human genomics. *Nature Methods*, pages 1–11, November 2024.
